# RNA Knowledge-Graph analysis through homogeneous embedding methods

**DOI:** 10.1101/2025.02.17.638592

**Authors:** Francesco Torgano, Mauricio Soto Gomez, Matteo Zignani, Jessica Gliozzo, Emanuele Cavalleri, Marco Mesiti, Elena Casiraghi, Giorgio Valentini

## Abstract

**Motivation:** We recently introduced RNA-KG, an ontology-based knowledge graph that integrates biological data on RNAs from over 60 public databases. RNA-KG captures functional relationships and interactions between RNA molecules and other biomolecules, chemicals, and biomedical concepts such as diseases and phenotypes, all represented within graph-structured bio-ontologies. We present the first comprehensive computational analysis of RNA-KG, evaluating the potential of graph representation learning and machine learning models to predict node types and edges within the graph.

**Results:** We performed node classification experiments to predict up to 81 distinct node types, and performed both generic and specific edge prediction tasks. Generic edge prediction focused on identifying the presence of an edge irrespective of its type, while specific edge prediction targeted specific interactions between ncRNAs, e.g. miRNA-miRNA or siRNA-mRNA, or relationships between ncRNA and biomedical concepts, e.g. miRNA-disease or lncRNA-Gene Ontology term relationships. Using embedding methods for homogeneous graphs, such as LINE and node2vec, in combination with machine learning models like decision trees and random forests, we achieved balanced accuracy exceeding 90% for the 20 most common node types and over 80% for most specific edge prediction tasks. These results show that simple embedding methods for homogeneous graphs can successfully predict nodes and edges of the RNA-KG, paving the way to discover novel ncRNA interactions and laying the foundation for further exploration and utilization of this rich information source to enhance prediction accuracy and support further research into the “RNA world”.

**Code Availability:** Python code to reproduce the experiments is available at https://github.com/AnacletoLAB/RNA-KG_homogeneous_emb_analysis

## 1. Introduction

Knowledge graphs (KGs) have been widely applied in biomedicine to collect and integrate diverse data types with concepts grounded in biomedical ontologies (Nicholson and Greene, 2020; Chandak et al., 2023).

We recently introduced RNA-KG, an ontology-based KG that integrates biological knowledge about coding and non-coding RNAs from more than 60 public databases (Cavalleri et al., 2024). RNA-KG incorporates functional relationships with genes, proteins, and chemicals, as well as ontologically grounded biomedical concepts. It was specifically designed to serve as input for AI-based techniques aimed at inferring novel knowledge about RNA molecules, thereby supporting RNA-drug design.

The structure and information encoded in KGs can be leveraged by graph representation learning (GRL) techniques to infer new knowledge. GRL encompasses a class of machine learning methods that encode graph-structured data as vectors, referred to as embeddings, while preserving the graph’s structural, relational, and attribute-based properties (Xia et al., 2021). These embeddings enable downstream tasks such as *link/edge prediction*, which identifies novel associations between concepts (nodes) in the graph, and *node/edge-type prediction*, which classifies node or edge types to reveal the structural organization of biological entities and their interactions. These techniques aid in identifying data patterns (Yi et al., 2021).

In this work, we present the first in-depth analysis of RNA-KG, employing homogeneous GRL techniques to determine whether relatively simple methods can effectively exploit the topological structure of local and global node neighborhoods to infer reliable knowledge from the KG and its biologically relevant subgraphs (views) extracted from RNA-KG.

More specifically, we first applied GRL methods to predict up to 81 distinct node types. Next, we performed *generic edge prediction*, which involves predicting the presence of any edge in the graph, irrespective of its type, where the edge type is determined by the types of the connected vertices. Finally, we employed GRL techniques for *specific edge prediction*, focusing on edges representing particular interactions between RNA molecules or relationships between RNA molecules and biomedical concepts. The motivation behind conducting both edge prediction experiments lies in their distinct objectives and practical applications. The generic edge prediction task evaluates whether the information encoded in the graph is sufficient and suitable for making accurate inferences, serving as a baseline assessment of the graph’s overall predictive power. However, in practical applications, the primary focus is often on determining whether specific nodes interact. For instance, one might seek to establish whether a particular siRNA (small interfering RNA) interacts with a target mRNA (messenger RNA) to induce RNA interference and knock down a specific gene (Damase et al., 2021). Similarly, specific miRNA-miRNA (microRNA) interactions are of interest, as a miRNA inhibitor could block the activity of another miRNA (Paunovska et al., 2022).

To address these practical needs, we introduced specific edge prediction tasks, each performed on specific RNA-KG views. These tasks enable more targeted predictions, such as inferring relationships between a miRNA and a disease by leveraging the disease ontologies included in RNA-KG.

## 2. Data and Methods

### 2.1. RNA-KG

RNA-KG (Cavalleri et al., 2024) is a Knowledge Graph (KG) that combines the publicly available information from more than 60 databases to obtain a centralized, uniform and semantically consistent representation of the “RNA world”. These molecules are widely studied because they have a primary role in biological processes and pathways, especially those that are altered in cancer, genetic disorders, neurodegenerative diseases, cardiovascular conditions, and infections (Damase et al., 2021). The study of RNA is also one of the most promising avenues of research in therapeutics, as evidenced by the recent success of mRNA-based vaccines for the COVID-19 pandemic (Barbier et al., 2022). RNA-KG represents the existing knowledge about interactions involving RNA molecules and their interactions with other biomolecular data as well as with chemicals, diseases, abnormal phenotypes, and proteins to support the study and the discovery of the biological role of the “RNA world”.

Supplementary table S1 provides detailed distributions of node types and edge types, highlighting the most prevalent categories in RNA-KG.

Among the three classification tasks described in Section 3, the tasks focused on specific-edge prediction were conducted on *RNA-KG views*. Each view was constructed to distill the information relevant to a classification task of particular importance in the biomedical domain and corresponds to a subgraph induced by node or edge types over the original RNA-KG.

We extracted seven views from RNA-KG, each tailored to encapsulate the biological information pertinent to a specific prediction task. These subgraphs were further enriched by integrating them with relevant portions of PheKnowLator’s Human Disease Benchmark KG (Callahan et al., 2024; Developers, 2024). This integration enabled the inclusion of additional biologically significant relationships, such as *gene-disease, disease-disease*, and *gene-gene* associations, thereby enhancing the contextual and predictive relevance of each view.

Figure 1 depicts the schema of the views, while in the supplementary section S2 we provide a description of each view, including node-type and edge-type distributions, as well as Complementary Cumulative Distribution Function (CCDF) plots for distinct node types within each view. These statistics offer a comprehensive overview of the structure and content of the views, highlighting their utility in addressing biologically relevant classification tasks, such as miRNA-disease or piRNA-disease prediction.

**Fig. 1:**
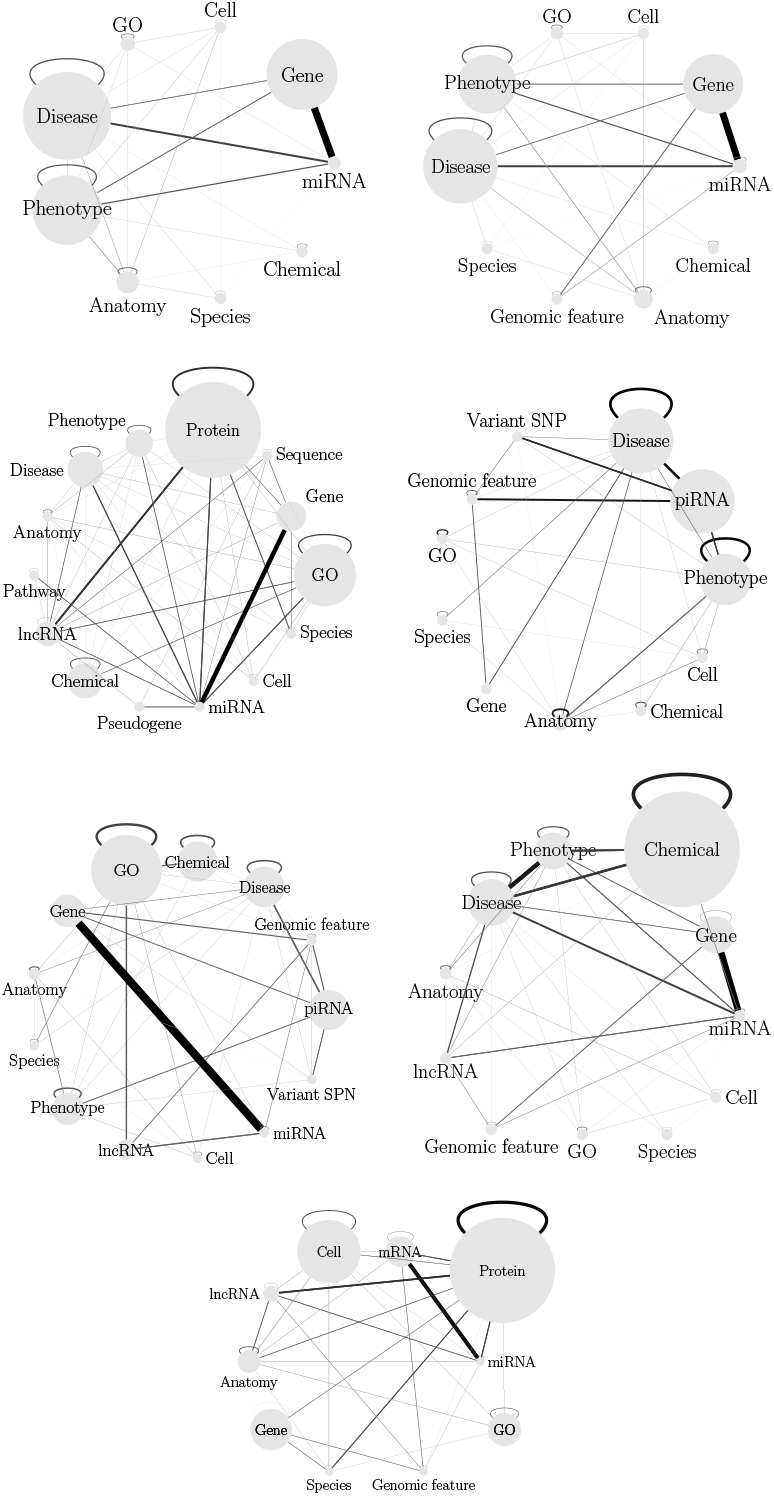
View schema. Each view is represented by a graph where node size is related to the number of nodes in the graph while edge width and color represent the proportion of edges between nodes of the respective types. In the figure we omitted node types accounting for less than 1% of the graph nodes.

### 2.2. Homogeneous Graph Representation Learning

Given a graph *G* = (*V, E*), where *V* is the set of nodes and *E* the set of edges, GRL techniques learn a function *f* : *G* → ℝ^*n*^ that maps (embeds) each node *v* ∈ *V* to a vector (alias, embedding) *x* ∈ ℝ^*n*^ such that nodes that are “close” in the graph *G* are also close in the vectorial space ℝ^*n*^. In other words, the embedding representation *X* of the node *v* preserves the topological characteristics of the node *v* in the embedding space ℝ^*n*^. These embeddings are then used by machine learning models to predict either the node/edge type or the existence of an edge.

Homogeneous GRL methods may be classified into the following three categories.

*Random walk-based methods* for homogeneous graphs like DeepWalk (Perozzi et al., 2014), Walklets (Perozzi et al., 2016) or node2vec (Grover and Leskovec, 2016) that use random walks on the graph to generate sequences of nodes that reflect the local and global connectivity patterns within the graph.

*Sampling-based methods* that try to preserve the proximity between the nodes, like LINE (Tang et al., 2015) that uses edge sampling methods to improve performance during the optimization of the objective function, designed to preserve both local and global graph structures.

*Graph Neural Network-based* methods like GraphSAGE (Hamilton et al., 2017) or Graph Attention Networks (Abu-El-Haija et al., 2018) can also be used to generate node embeddings. Graph Neural Networks showed excellent performance in graph prediction tasks, but require more complex and computationally intensive models (Ma et al., 2022).

Although homogeneous GRL methods simplify the embedding process by neglecting node and edge type information, they have proven their effectiveness and efficiency in the information inference from graph-structured data. In particular, homogeneous GRL techniques are less computationally demanding than heterogeneous methods, making them more suitable for large-scale graphs. Moreover, they avoid issues such as type imbalance that can arise in heterogeneous techniques, where the most represented types dominate the embeddings.

In this work, we investigate whether simple embedding methods can infer reliable knowledge from the RNA-KG or from smaller, biologically relevant, subgraphs (views) extracted from the overall graph. The two homogeneous embedding methods used in this study, node2vec (a random-walk-based method) and LINE (a sampling-based technique), were carefully selected based on their computational efficiency, simplicity, and proven performance. LINE was chosen for its speed, which allows for rapid experimentation and testing, while node2vec was selected for its demonstrated effectiveness in diverse graph-prediction tasks. The embeddings by these methods are independent of the subsequent classification task, which can be performed using the embeddings as input to any supervised machine learning model, even as relatively simple as those we tested in our study: DTs (Breiman et al., 1984), RFs (Breiman, 2001)^1^.

## 3. Results

In this section, we describe the results obtained by embedding the graph elements with LINE and node2vec (under the settings defined in subsection 3.1), and then training DTs, and RFs on such embeddings for three classification tasks: 1. Node-type prediction tasks applied to the whole RNA-KG (subsection 3.2); 2. Generic edge prediction tasks applied to the whole RNA-KG (Subsection 3.4); 3. Specific edge prediction tasks, each applied to a specific RNA-KG view (Subsection 3.5).

### 3.1. Graph embedding methods

To embed the graph with LINE, we utilized both the first-order and second-order proximity versions provided by the GRAPE library (Cappelletti et al., 2023), using the default hyperparameters.

For node2vec, we adopted the skip-gram model as the shallow neural network architecture, with GRAPE default hyperparameters.

To ensure a fair comparison, we generated random walk samples across multiple experiments using the same setting, defined by the same values for the `return_weight` 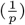 and `explore_weight` 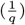 hyperparameters. More precisely we compared values that emulate a depth-first visit (*DFS* -setting with 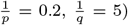, a breadth-first visit (*BFS* -setting with 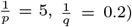, or an unbiased first-order random walk according to a DeepWalk-like strategy (Perozzi et al., 2014) (*Balanced* -setting with 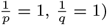.

Figure 2 shows the t-SNE two-dimensional projections (van der Maaten and Hinton, 2008) of the LINE and node2vec embeddings for the most frequent types of nodes in RNA-KG. Our preliminary results showed that the BFS-like node2vec strategy performs better in the predictive tasks. This strategy has been used in all the experiments reported in section 3.

**Fig. 2:**
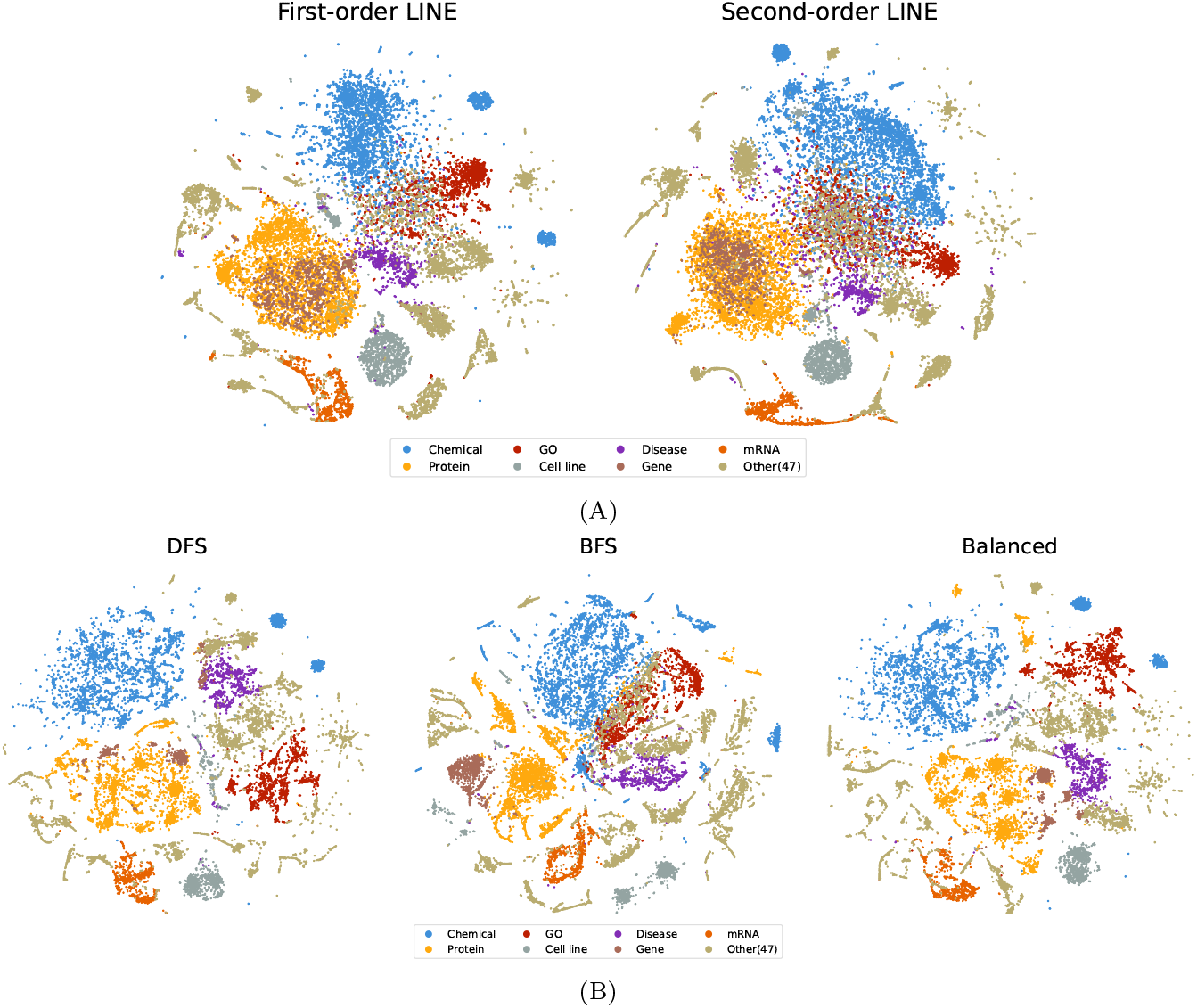
2D t-SNE projections of node embeddings of the RNA-KG. (A) First (left) and second-order (right) LINE node embeddings; (B) node2vec with DFS-like (left) and BFS-like (center) hyperparameters and DeepWalk-like (right) node embeddings.

### 3.2. Node-type prediction: experimental settings and results

For node-type prediction we computed embeddings by using BFS-like node2vec to project points into multiple embedding sizes: 10, 50 and 100, and we also experimented with 2-dimensional projections of 100-dimensional embeddings through t-SNE. On average, 10-D embeddings achieved the best balance between predictive accuracy and efficiency (more details in Supplementary section S3). The embeddings were input to DTs and RFs. To perform an unbiased evaluation, we applied 5 stratified holdout cross-validation (train:test ratio = 70:30, with fixed seed for the random number generator to guarantee comparable results across models) on the RNA-KG nodes (about 578K nodes). Hyperparameter selection for the classifier models was performed on the training set using a grid search strategy and internal cross-validation.

Figure 3 shows the average balanced accuracy results obtained from the 10-D embeddings. Overall results are surprisingly positive. Using RFs, we achieve a balanced accuracy of 97.4% when classifying the 7 most represented node types. Even when considering the 20 main node types, the results remain well above 90%. When considering 81 different node types, the balanced accuracy drops to 53.7%; however, it is important to note that a random guess predictor would achieve a balanced accuracy of only about 1.2%. Details about the considered node types are available in the Supplementary Section S1.

**Fig. 3:**
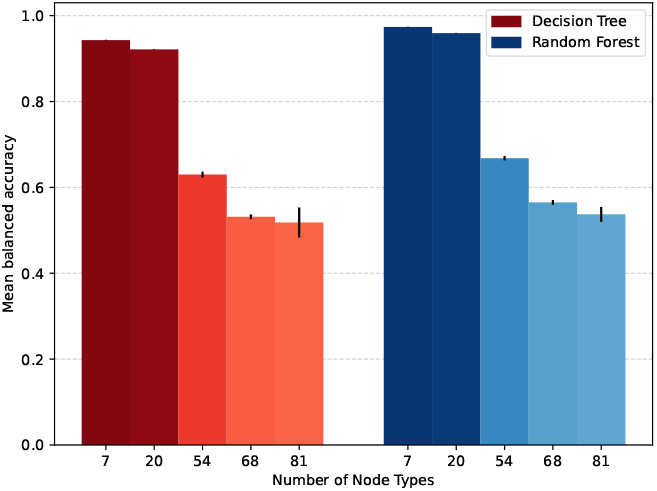
Balanced accuracy results on RNA-KG node type prediction using full 10-D BFS-like node2vec embeddings with DTs (red bars) and RFs (blue bars), considering the 7, 20, 54, 68 and 81 most common node types. The color intensity of the bars reflects their balanced accuracy value. Vertical segments on each bar represent the standard deviation across multiple holdouts; note that in some cases the standard deviation is very low (*<* 0.1%) and the segment is not visible.

### 3.3. Edge prediction: unbiased sampling of train and test sets

We experimented with generic and specific edge prediction tasks. Fig. 4 depicts the main differences between these two types of edge prediction.

**Fig. 4:**
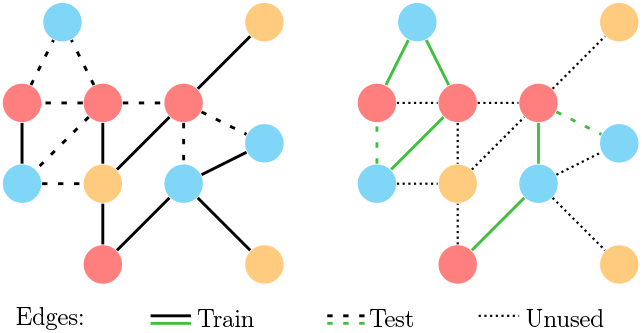
Generic (left) and specific (right) edge prediction, where the specific edges of interest are the green ones. Continuous lines represent positive training edges, dashed ones positive test edges; dotted lines in the right panel represent the edges not used for either training or testing in the specific edge prediction.

When performing edge prediction tasks, a commonly overlooked issue is the method used to create training and test edge samples.

The first important observation is that, since test edges must be removed from the input graph prior to graph embedding, their removal should not alter the graph’s global connectivity structure; specifically, we should not change the number of disconnected components. To ensure this, for both generic and specific edge prediction tasks, we generated training and test splits using a *Connected Monte-Carlo* holdout scheme, which ensures that removing test edges does not increase the number of disconnected components compared to the original full RNA-KG (Cappelletti et al., 2023).

In the context of specific edge prediction, particular attention should be given to the generation of negative edges, to avoid false negatives that can compromise the learning capabilities of the models. Indeed, if the graph used to generate the negative training edges is limited to the positive training graph, rather than the full graph, positive examples that are not included in the training set but that have been included in the test set could lead to false negatives that can confuse the predictor being trained, degrading its performance. This procedure, included in GRAPE, can be acceptable when we consider generic edge prediction of the overall graph since the expected number of false negatives is relatively low, but with specific edges, the number of potential false negatives can be large, as we will show in the following experiments.

For this reason, we designed and implemented the following unbiased pipeline for specific edge prediction:

1. Generate the positive train 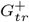 and positive test 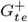 graph from the full graph *G*, while guaranteeing the connectivity and the same number of connected components in the positive train graph 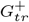 as in the original graph *G*. This can be accomplished by using a spanning tree of *G* as 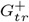, and 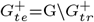 .
2. Generate the embeddings for the nodes of the graph by applying an embedding method on the training graph 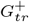 (the embeddings are generated based on the train graph instead of the full graph to avoid bias in the results).
3. Filter the positive train 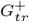 and positive test graph 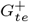 edges by keeping only the specific edge type to obtain a filtered positive train 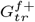 and test graph 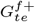.
4. Train the edge prediction model using a novel, unbiased training procedure that uses the full graph *G* to generate the negative training graph 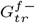, instead of the filtered train graph 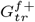 utilized by GRAPE. This substitution was made because using the positive filtered train graph 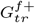 to generate the negative training graph 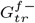 can lead to a negative graph that contains false negatives.
5. Generate the negative test graph 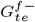 from the full graph *G* (not from the hold-out, to avoid positive edges being extracted as negative) with the source and destination type we are interested in, using node-degree aware edge sampling to guarantee that the positive and negative edges included have comparable degree distributions (Cappelletti et al., 2024).
6. Test the trained model with the filtered positive test graph 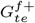 and the negative test graph 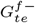.

Details about the implementation of the unbiased train/test sampling approach can be found in Supplementary Section S6.

### 3.4. Generic Edge Prediction

Generic edge prediction experiments have been performed using 5 stratified holdout cross-validation, with a 70:30 train:test ratio, and using the same random seed generator to guarantee comparable results across models. Overall, the training set contains about 4M edges, and the test set with about 1.7M edges.

Both DTs and RFs hyperparameters are optimized by using grid-search on the training set (learning hyperparameters listed in Supplementary section S4).

Experimental results with different embedding dimensions are shown in Figure 5. The embedding size has a certain impact on the Balanced Accuracy only with RFs. The best result is achieved with RFs trained on 100-D embeddings. Again, results are encouraging and confirm the predictive capability of homogeneous embedding methods on RNA-KG.

**Fig. 5:**
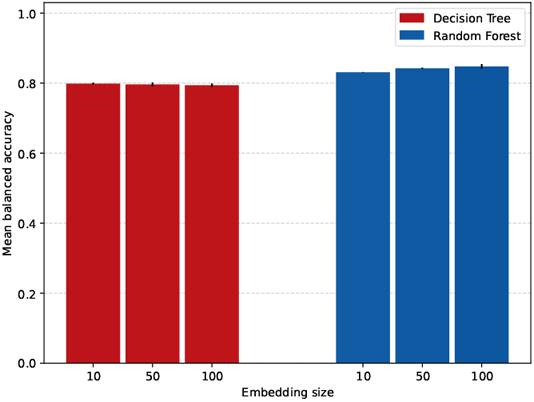
Balanced accuracy results for DT (red bars) and RF (blue bars) on the RNA-KG dataset for the generic edge prediction task, utilizing full BFS-like node2vec embeddings of various sizes. Vertical segments on each bar represent the standard deviation across multiple holdouts; note that in some cases the standard deviation is very low (*<* 0.1%) and the segment is not visible.

### 3.5. Specific Edge Prediction

For the specific edge prediction tasks, we embedded each RNA-KG view by three embedding approaches, i.e., BFS and DFS-like node2vec Skip-Gram and first-order LINE, and trained RFs and DTs to classify the computed embeddings (all the experiments employed the same hyperparameter values, see Supplementary section S5).

Generalization capabilities of the models were assessed using 5 random holdout cross-validation (train:test ratio = 70:30, with random seed generator set to a fixed value) in all the following experiments.

#### 3.5.1. Specific edge prediction results with biased/unbiased train/test sampling

We compared the biased and unbiased train/test sampling approaches (Section 3.3) on a subset of RNA-KG views. The embeddings were generated using first-order LINE with embedding size set to 10 and the DT as classifier model.

Table 1 shows that the biased sampling can generate a consistent percentage of false negative edges (FN% column) in some specific edge prediction tasks (e.g. 21% and 23% in, respectively, the miRNA-Disease and miRNA-Phenotype tasks). The performance of the classifier models are therefore consequently affected, as witnessed by the accuracy on the positive edges that drops to 50%. This issue is solved when using the model trained with the unbiased training pipeline; indeed, the accuracy on the positive edges improves to more than 75% in both cases. On the other hand, results on negative edges are not affected, thus resulting in boosted balanced accuracy. As expected, when the percentage of false negatives is low, the performance obtained when using the biased and unbiased sampling approaches are comparable (Table 1).

**Table 1.** Comparison of the biased and unbiased pipeline on the miRNA-disease view of the RNA-KG across six edge specific prediction tasks. FN stands for False Negatives and BA for Balanced Accuracy.

| Specific edge prediction | Unbiased Procedure | FN(%) | Accuracy on pos. | Accuracy on neg. | Overall BA(%) |
| --- | --- | --- | --- | --- | --- |
| miRNA-Disease | No | 21.27 | 50.10 ± 0.21 | 75.31 ± 0.18 | 62.71 ± 0.11 |
| miRNA-Disease | Yes | 0.00 | 75.15 ± 0.34 | 76.99 ± 0.31 | 76.07 ± 0.14 |
| Disease-Disease | No | 0.17 | 48.34 ± 0.53 | 63.48 ± 0.29 | 55.91 ± 0.23 |
| Disease-Disease | Yes | 0.00 | 47.18 ± 0.61 | 64.01 ± 0.96 | 55.60 ± 0.39 |
| miRNA-Gene | No | 4.00 | 56.03 ± 0.12 | 59.14 ± 0.11 | 57.59 ± 0.08 |
| miRNA-Gene | Yes | 0.00 | 59.39 ± 0.11 | 59.36 ± 0.08 | 59.38 ± 0.09 |
| Gene-Disease | No | 0.51 | 52.39 ± 0.83 | 62.63 ± 0.75 | 57.51 ± 0.34 |
| Gene-Disease | Yes | 0.00 | 51.70 ± 0.61 | 63.14 ± 0.43 | 57.42 ± 0.32 |
| Gene-Phenotype | No | 1.26 | 53.55 ± 0.59 | 54.57 ± 0.55 | 54.06 ± 0.35 |
| Gene-Phenotype | Yes | 0.00 | 53.65 ± 0.48 | 55.16 ± 0.96 | 54.41 ± 0.57 |
| miRNA-Phenotype | No | 23.15 | 50.25 ± 0.36 | 77.20 ± 0.32 | 63.73 ± 0.17 |
| miRNA-Phenotype | Yes | 0.00 | 75.72 ± 0.63 | 78.42 ± 0.39 | 77.07 ± 0.29 |

#### 3.5.2. Specific edge prediction results with the unbiased pipeline on different views of the RNA-KG

Finally, we conducted a thorough analysis of various views of the RNA-KG, each extracted from the overall RNA-KG to focus on specific types of nodes and edges relevant to a given edge prediction task.

The balanced accuracy results for the main specific edge prediction tasks are summarized in Fig. 6. These results were obtained using a 10-D LINE embedding combined with DTs and RFs.

**Fig. 6:**
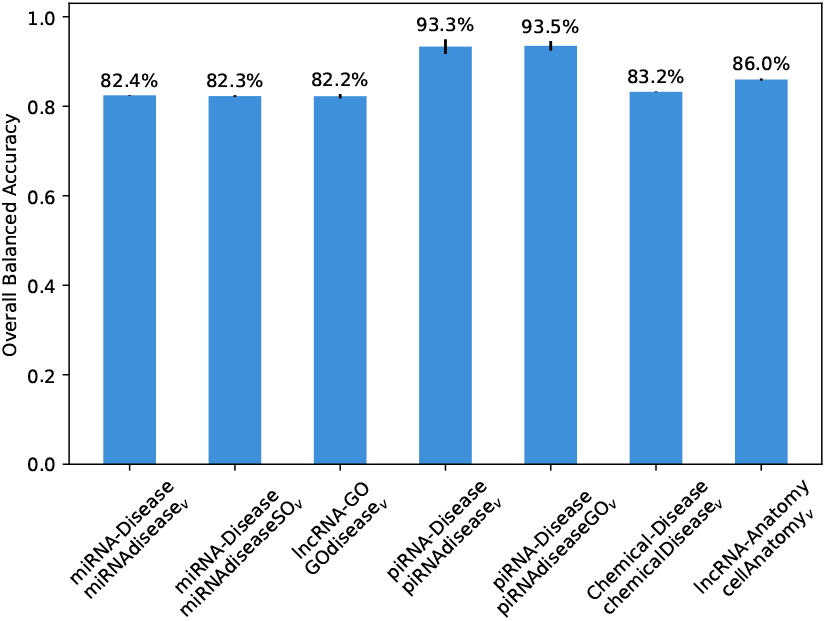
Comparison of the balanced accuracy achieved by the RFs on specific edge prediction tasks of seven RNA-KG views. The first line on the x axis refers to the specific edge prediction task, the second line to the RNA-KG view. The vertical line on top of the bars represents the standard deviation of an holdout procedure repeated 5 times.

Detailed results for multiple specific edge prediction tasks across seven RNA-KG views are provided in Supplementary section S5.

## 4. Discussion

In this work, we conducted thorough and unbiased experiments on RNA-KG. Our results demonstrate that embedding methods designed for homogeneous graphs, such as node2vec, achieve surprisingly good performance even when applied to a fully heterogeneous graph like RNA-KG, which is characterized by a large number of node and edge types.

Indeed, as shown in Fig. 3, the node-type prediction results yield very high balanced accuracy using relatively simple methods (e.g., DTs), although the best performance is achieved with RFs. Full embeddings lead to significantly better results (Wilcoxon rank sum test, *p*-value *<* 10^−5^), which is expected since 2D projections overly compress the data and result in a loss of important features.

Additionally, for the generic edge prediction task, RFs trained on embedded edges achieve accuracy greater than 80%, confirming that embedding methods for homogeneous graphs can reasonably predict the existence of an edge in RNA-KG.

Edge prediction tasks focusing on specific edges in RNA-KG, using views tailored to the prediction task, show that we can obtain accuracy larger than 80%. In several cases, such as piRNA-disease prediction, accuracy exceeds 90% (Fig. 6). This performance is achieved using relatively simple and fast embedding methods, such as LINE, in combination with off-the-shelf classifiers like DTs and RFs.

Overall, these results indicate that we can reasonably predict nodes and edges in RNA-KG using relatively simple embedding methods for homogeneous graphs, thereby demonstrating both the effectiveness of these methods and the high quality of the data available in RNA-KG. This is of paramount importance for discovering novel interactions between different types of non-coding RNA or other biomolecular entities, or for uncovering relationships between RNAs and specific biomedical concepts (e.g., diseases or phenotypes).

These findings represent the first systematic analysis of RNA-KG using relatively simple embedding methods for homogeneous graphs.

We estimate that the success of these methods partially relies on the topological differences between node types. Indeed, as shown in Table S1 in the supplementary information, all views exhibit a substantial variability in the mean degree across node types. This property could facilitate a characterization of node and edge types based on their neighborhoods, which can be exploited by homogeneous methods. These differences are even more pronounced when considering the semantic induced of node types. Therefore, we anticipate substantial improvements in prediction performance by employing methods that account for the rich semantic heterogeneity of the graph (Bing et al., 2023), thereby exploiting the available knowledge about the diverse types of nodes and edges in RNA-KG (Soto-Gomez et al., 2024). Furthermore, considering that RNA-KG is constructed by integrating ontologies that hierarchically organize information, we plan to explore hyperbolic embedding techniques, as hyperbolic spaces are known to better model hierarchical information compared to Euclidean spaces (Nickel and Kiela (2017); Sala et al. (2018)).

In summary, our results show that GRL methods can accurately predict node types and edges in RNA-KG. The resulting list of predicted interactions and relationships could serve as an invaluable resource for guiding the experimental efforts of biomedical researchers, thus paving the way for novel discoveries and insights into the “RNA-world”.

## 5. Competing interests

No competing interest is declared.

## 6. Author contributions statement

FT and MSG: performed visualizations; MSG, ElC, and GV: designed the study; FT, MSG: developed the software; FT, MSG, EmC: curated the data; FT, MSG: performed experiments; FT, MSG, MM, MZ, ElC, GV: validated results; FT, MSG, JG, ElC, GV: wrote the original draft; MSG, JG, MM, MZ, ElC, GV: reviewed the original draft; ElC, GV: acquired fundings, supervised the work, conceptualized the study. All the authors read and approved the manuscript.

## 7. Code and data availability

RNA-KG is available at https://rna-kg.anacleto.di.unimi.it/ and RNA-KG views are available at https://rna-kg.anacleto.di.unimi.it/views/. The RNA-KG version used is 0.2 which is also available to download from https://zenodo.org/records/10078877. The code that was used to run the experiments is available on GitHub (https://github.com/AnacletoLAB/RNA-KG_homogeneous_emb_analysis).

## 8. Acknowledgments

This work is supported by National Center for Gene Therapy and Drugs Based on RNA Technology—MUR (Project no. CN 00000041) funded by NextGeneration EU program.

## Footnotes

1 Note that we also tested radial basis function support vector machines (RBF-SVMs), which achieved results similar to RFs. Due to the superior robustness of RFs w.r.t. their hyperparameter settings and their fastest convergence, we preferred reporting RF results. All the models were implemented using Python - *scikit-learn* (Pedregosa et al., 2011) library.

## Notes

https://github.com/AnacletoLAB/RNA-KG_homogeneous_emb_analysis

### Competing Interest Statement

The authors have declared no competing interest.

