## Supplementary_material for "RNA Knowledge-Graph analysis through homogeneous embedding methods"

**S3 Node Type Prediction** **19**

**S4 Generic Edge Prediction** **20**

**S5 Specific Edge Prediction** **23**

#### S1 RNA-KG node types

Supplementary Table S1 lists all the node types available in RNA-KG along with their respective counts.

### S2 RNA-KG views

In this section, we provide a detailed description of the different views used for the predictive tasks.

To describe the main structure of the views and highlight their differences, for

Table S1: Node distribution respect to the node type for RNA-KG.

| Node | Count | Proportion | Mean degree |
|---|---|---|---|
| Chemical | 150080 | 0.2595 | 4.1148 |
| Protein | 96197 | 0.1663 | 11.1001 |
| GO | 43869 | 0.0758 | 11.2065 |
| Cell line | 41807 | 0.0723 | 7.0619 |
| Disease | 23262 | 0.0402 | 7.9286 |
| tRF | 20897 | 0.0361 | 13.2522 |
| mRNA | 20624 | 0.0357 | 121.1773 |
| Gene | 20081 | 0.0347 | 4.9014 |
| Phenotype | 16865 | 0.0292 | 6.3985 |
| Variant (SNP) | 16091 | 0.0278 | 3.0458 |
| Riboswitch | 15805 | 0.0273 | 8.7789 |
| lncRNA | 15511 | 0.0268 | 92.1723 |
| miRNA | 14295 | 0.0247 | 145.3281 |
| Anatomy | 14203 | 0.0246 | 7.8024 |
| tsRNA | 10243 | 0.0177 | 15.3638 |
| Pseudogene | 9837 | 0.0170 | 27.1647 |
| circRNA | 9216 | 0.0159 | 48.0559 |
| Small protein | 8178 | 0.0141 | 3.0584 |
| Vaccine | 6246 | 0.0108 | 4.9922 |
| Viral RNA | 5654 | 0.0098 | 2.0428 |
| ASO | 2633 | 0.0046 | 2.0171 |
| Pathway | 2600 | 0.0045 | 2.5454 |
| Cell | 2368 | 0.0041 | 18.2247 |
| Sequence | 2363 | 0.0041 | 78.6022 |
| Species | 2148 | 0.0037 | 60.1378 |
| snRNA | 1803 | 0.0031 | 16.7598 |
| tRNA | 894 | 0.0015 | 258.6521 |
| Glycan | 797 | 0.0014 | 11.7465 |
| snoRNA | 583 | 0.0010 | 31.2727 |
| otherRNA | 565 | 0.0010 | 22.8053 |
| Environment | 398 | 0.0007 | 4.6533 |
| rRNA | 350 | 0.0006 | 5.4857 |
| Chromosome | 249 | 0.0004 | 4.1606 |
| Environment exposure | 171 | 0.0003 | 4.7310 |
| Neuro behaviour | 165 | 0.0003 | 2.6667 |
| Food | 163 | 0.0003 | 3.1288 |
| Experimenta factor | 145 | 0.0003 | 73.4138 |
| Aptamer | 138 | 0.0002 | 2.1594 |
| s(i/h)RNA | 117 | 0.0002 | 2.0342 |
| ncRNA | 115 | 0.0002 | 10.8522 |
| Protein modification | 91 | 0.0002 | 117.9011 |
| gRNA | 77 | 0.0001 | 2.0000 |
| Mouse pathology | 75 | 0.0001 | 5.6000 |
| Medical action | 72 | 0.0001 | 1.9861 |
| Plant | 48 | 0.0001 | 3.8542 |
| viral_mRNA | 39 | 0.0001 | 7.4359 |
| NCI thesaurus | 28 | 0.0000 | 3.0714 |
| scaRNA | 23 | 0.0000 | 24.4783 |
| lincRNA | 22 | 0.0000 | 3.5455 |
| Histone modification | 20 | 0.0000 | 3675.3000 |
| Epigenetic modification | 18 | 0.0000 | 13.7778 |
| Human developmental stage | 16 | 0.0000 | 4.9375 |
| Mental functioning | 14 | 0.0000 | 2.8571 |
| RNA drug | 13 | 0.0000 | 4.4615 |
| mtRNA | 10 | 0.0000 | 4.8000 |
| retained_intron | 8 | 0.0000 | 5.1250 |
| unknown RNA | 8 | 0.0000 | 14.6250 |
| undefined | 6 | 0.0000 | 17.5000 |
| General medical science | 6 | 0.0000 | 2.3333 |
| CP | 6 | 0.0000 | 2.0000 |
| Mental disease | 6 | 0.0000 | 2.0000 |
| PCO | 6 | 0.0000 | 3.1667 |
| Ribozyme | 6 | 0.0000 | 983.6667 |
| Y_RNA | 4 | 0.0000 | 3.0000 |
| Adverse events | 4 | 0.0000 | 1.7500 |
| eRNA | 3 | 0.0000 | 2.6667 |
| vRNA | 3 | 0.0000 | 5.0000 |
| ribozyme | 3 | 0.0000 | 36.6667 |
| ExO | 3 | 0.0000 | 61.3333 |
| scRNA | 3 | 0.0000 | 36.0000 |
| Biological role | 3 | 0.0000 | 57.0000 |
| snomedct | 2 | 0.0000 | 10938.5000 |
| Infectious disease | 2 | 0.0000 | 1.0000 |
| misc_RNA | 2 | 0.0000 | 2.0000 |
| TEC | 2 | 0.0000 | 2.0000 |
| Basic formal | 1 | 0.0000 | 1113.0000 |
| Trait | 1 | 0.0000 | 2.0000 |
| UMLS | 1 | 0.0000 | 2.0000 |
| Drosophila anatomy | 1 | 0.0000 | 2.0000 |
| piRNA | 1 | 0.0000 | 2.0000 |
| owlNothing | 1 | 0.0000 | 208.0000 |

Figure S1: RNA-KG schema. In the graph, node size represent the proportion of nodes in the graph while edge width and color represent the proportion of edges between nodes of the respective types. In the figure we omitted node types accounting for less than 0.15% of the graph nodes.

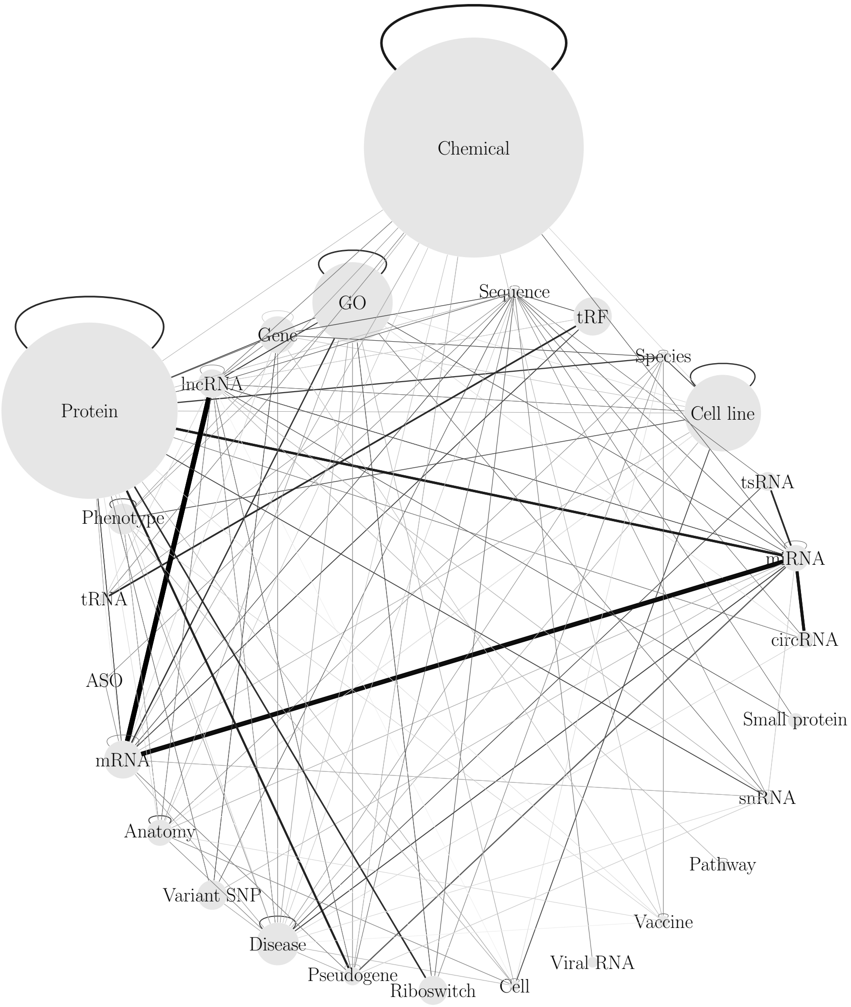

each of them we plot both the complementary cumulative distribution function for the node degrees with respect to the node types and the graph schema. Additionally, we describe the distribution of the node types and of the edge types.

In figure S2, we represent the complementary cumulative distribution function (CCDF) of all the RNA-KG views.

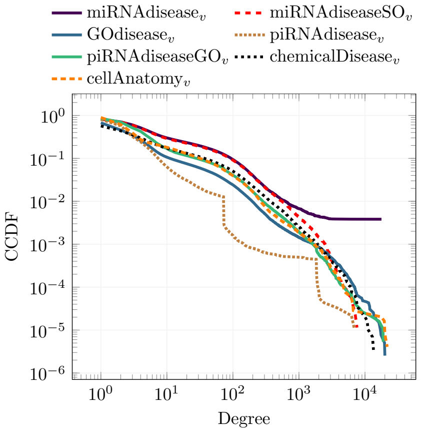

Figure S2: Complementary Cumulative Degree Distribution (CCDF) of RNA-KG views.

#### S2.1 miRNAdisease

Figure S3: Left: Complementary Cumulative Distribution Function (CCDF) of the node degrees by node type for the miRNAdisease view. Right: Graph induced by the view types. In the graph, node size represent the proportion of nodes in the graph while edge width and color represent the proportion of edges between nodes of the respective types. In the figure we omitted node types accounting for less than 1% of the graph nodes.

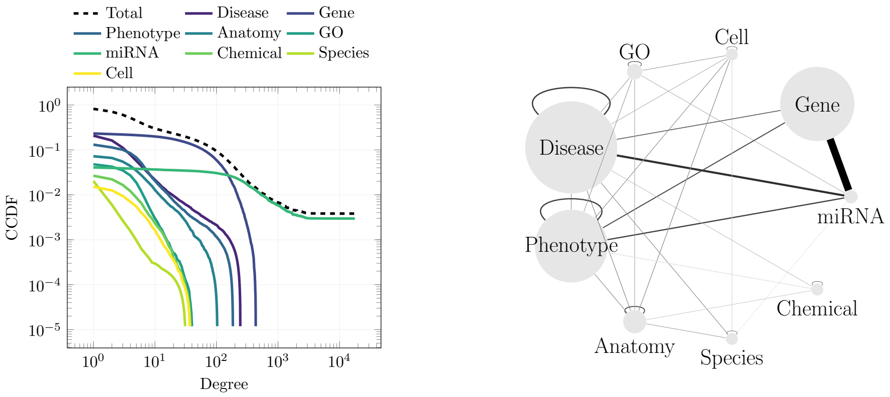

Table S2: Node distribution respect to the node type for the miRNAdisease view*.

| **Node** | **Count** | **Proportion** | **Mean degree** |
|---|---|---|---|
| Disease | 24203 | 0.2882 | 9.0892 |
| Gene | 19517 | 0.2324 | 71.7402 |
| Phenotype | 19152 | 0.2280 | 7.7664 |
| Anatomy | 6084 | 0.0724 | 8.6917 |
| GO | 4050 | 0.0482 | 5.9427 |
| miRNA | 3493 | 0.0416 | 423.4423 |
| Chemical | 2239 | 0.0267 | 5.6829 |
| Species | 1784 | 0.0212 | 3.1508 |
| Cell | 1272 | 0.0151 | 5.9041 |
| Environment | 453 | 0.0054 | 4.8234 |
| Environmental exposure | 419 | 0.0050 | 3.7613 |
| Chromosome | 310 | 0.0037 | 5.1839 |
| Protein | 205 | 0.0024 | 7.3366 |
| Food | 181 | 0.0022 | 2.8619 |
| Neuro behaviour | 153 | 0.0018 | 3.7647 |
| Medical action | 114 | 0.0014 | 3.7895 |
| Genomic feature | 98 | 0.0012 | 276.2857 |
| NCI thesaurus | 84 | 0.0010 | 3.1310 |

* We report only node types accounting for more than 0.1% of the graph nodes.

Table S3: Edge distribution respect to the type of the edge node extremes for the miRNAdisease view. We include the most frequent predicates for the extreme node types*.

| **Source** | **Destination** | **Count** | **Proportion** | **Predicates** |
|---|---|---|---|---|
| miRNA | Gene | 1333269 | 0.7869 | Involved in regulation of |
| miRNA | Disease | 105211 | 0.0621 | Causes or contributes to condition |
| Disease | Disease | 44117 | 0.0260 | Subclassof (0.9444), Has characteristic (0.0367), Predisposes towards (0.0085) |
| Phenotype | Phenotype | 38489 | 0.0227 | Subclassof (25475/0.6619), Has modifier (0.1672), Has quality (0.1563) |
| miRNA | Phenotype | 36752 | 0.0217 | Causes or contributes to condition |
| Gene | Phenotype | 24519 | 0.0145 | Causes or contributes to condition |
| Gene | Genomic feature | 23322 | 0.0138 | Subclassof |
| Anatomy | Anatomy | 20840 | 0.0123 | Subclassof (0.5189), Part of (0.2536), Develops from (0.0452), Connects (0.0167), Has part (0.0139), Composed primarily of (0.0138), Contributes to morphology of (0.0118) |
| Gene | Disease | 14422 | 0.0085 | Causes or contributes to condition |
| GO | GO | 10978 | 0.0065 | Subclassof (0.7719), Part of (0.0580), Regulates (0.0575), Positively regulates (0.0505), Negatively regulates (0.0504) |
| Phenotype | Anatomy | 6479 | 0.0038 | Characteristic of (0.5309), Part of (0.2423), Characteristic of part of (0.1375) |
| Chemical | Chemical | 5844 | 0.0034 | Subclassof (0.5643), Has role (0.2033) |
| Disease | Gene | 4622 | 0.0027 | Has material basis in germline mutation in (0.9939) |
| miRNA | Genomic feature | 3493 | 0.0021 | Subclassof |
| Disease | Anatomy | 2996 | 0.0018 | Disease has location (0.8872), Disease has inflammation site (0.0844), Disease arises from alteration in structure (0.0197) |
| Cell | Cell | 2380 | 0.0014 | Subclassof (0.8924), Develops from (0.0983) |
| Species | Species | 1782 | 0.0011 | Subclassof |
| Disease | Phenotype | 1655 | 0.0010 | Disease has feature (0.4961), Has characteristic (0.3994), Disease has major feature (0.0550), Disease arises from feature (0.0387) |

* We report only pair of node types accounting for more than 0.1% of the graph edges; and for each pair of node types, the predicates accounting for more than 1% of the edge set.

#### S2.2 miRNAdiseaseSO

Figure S4: Left: Complementary Cumulative Distribution Function (CCDF) of the node degrees by node type for the miRNAdiseaseSO view. Right: Graph induced by the view types. In the graph, node size represent the proportion of nodes in the graph while edge width and color represent the proportion of edges between nodes of the respective types. In the figure we omitted node types accounting for less than 1% of the graph nodes.

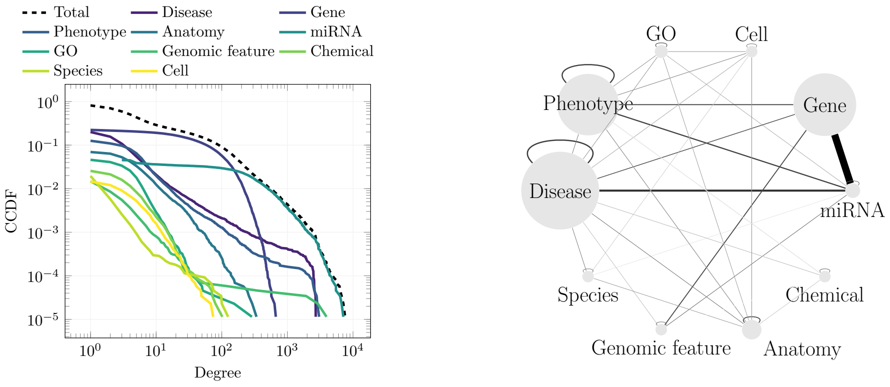

Table S4: Node distribution respect to the node type for the miRNAdiseaseSO view*.

| **Node** | **Count** | **Proportion** | **Mean degree** |
|---|---|---|---|
| Disease | 24203 | 0.2771 | 9.0895 |
| Gene | 19517 | 0.2234 | 71.7402 |
| Phenotype | 19152 | 0.2192 | 7.7664 |
| Anatomy | 6084 | 0.0696 | 8.6915 |
| miRNA | 4565 | 0.0523 | 326.7593 |
| GO | 4050 | 0.0464 | 5.9407 |
| Genomic feature | 2391 | 0.0274 | 14.3141 |
| Chemical | 2239 | 0.0256 | 5.6829 |
| Species | 1784 | 0.0204 | 3.1508 |
| Cell | 1272 | 0.0146 | 5.9041 |
| Environment | 453 | 0.0052 | 4.8256 |
| Environmental exposure | 419 | 0.0048 | 3.7613 |
| Chromosome | 310 | 0.0035 | 5.1839 |
| Protein | 205 | 0.0023 | 7.3366 |
| Food | 181 | 0.0021 | 2.8674 |
| Neuro behaviour | 153 | 0.0018 | 3.7647 |
| Medical action | 114 | 0.0013 | 3.7895 |
| NCI thesaurus | 84 | 0.0010 | 3.1190 |

* We report only node types accounting for more than 0.1% of the graph nodes.

Table S5: Edge distribution respect to the type of the edge node extremes for the miRNAdiseasSO view. We include the most frequent predicates for the extreme node types*.

| **Source** | **Destination** | **Count** | **Proportion** | **Predicates** |
|---|---|---|---|---|
| miRNA | Gene | 1333269 | 0.7824 | Involved in regulation of |
| miRNA | Disease | 105211 | 0.0617 | Causes or contributes to condition |
| Disease | Disease | 44117 | 0.0259 | Subclassof (0.9444), Has characteristic (0.0367) |
| Phenotype | Phenotype | 38489 | 0.0226 | Subclassof (0.6619), Has modifier (0.1672), Has quality (0.1563), Has part (0.0104), Increased in magnitude relative to (0.0014) |
| miRNA | Phenotype | 36752 | 0.0216 | Causes or contributes to condition |
| Gene | Phenotype | 24519 | 0.0144 | Causes or contributes to condition |
| Gene | Genomic feat. | 23322 | 0.0137 | Subclassof |
| Anatomy | Anatomy | 20840 | 0.0122 | Subclassof (0.5189), Part of (0.2491), Develops from (0.0457), Connects (0.0167), Contributes to morphology of (0.0143), Has part (0.0139), Composed primarily of (0.0138) |
| Gene | Disease | 14422 | 0.0085 | Causes or contributes to condition |
| GO | GO | 10978 | 0.0064 | Subclassof (0.7719), Part of (0.0580), Regulates (0.0575), Positively regulates (0.0505), Negatively regulates (0.0504) |
| Phenotype | Anatomy | 6479 | 0.0038 | Characteristic of (0.5314), Part of (0.2423), Characteristic of part of (0.1375), Occurs in (0.0585), Towards (0.0252) |
| Chemical | Chemical | 5844 | 0.0034 | Subclassof (0.5643), Has role (0.2033), Is conjugate acid of (0.0647), Is conjugate base of (0.0647), Has functional parent (0.0320), Is tautomer of (0.0250), Is enantiomer of (0.0192), Has part (0.0171) |
| miRNA | miRNA | 5750 | 0.0034 | Develops from (0.5000), Develops into (0.5000), |
| Disease | Gene | 4622 | 0.0027 | Has material basis in germline mutation in (0.9939) |
| miRNA | Genomic feat. | 4565 | 0.0027 | Subclassof |
| Genomic feat. | Genomic feat. | 3145 | 0.0018 | Subclassof (0.8003), Part of (0.0585), Has quality (0.0525), Has part (0.0483), Derives from (0.0207) |
| Disease | Anatomy | 2996 | 0.0018 | Disease has location (0.8868), Disease has inflammation site (0.0848), Disease arises from alteration in structure (0.0197) |
| Cell | Cell | 2380 | 0.0014 | Subclassof (0.8924), Develops from (0.0983) |
| Species | Species | 1782 | 0.0010 | Subclassof |
| Disease | Phenotype | 1655 | 0.0010 | Disease has feature (0.4967), Has characteristic (0.3994), Disease has major feature (0.0550), Disease arises from feature (0.0381) |

* We report only pair of node types accounting for more than 0.1% of the graph edges; and for each pair of node types, the predicates accounting for more than 1% of the edge set.

#### S2.3 GOdisease

Figure S5: Left: Complementary Cumulative Distribution Function (CCDF) of the node degrees by node type for the GOdisease view. Right: Graph induced by the view types. In the graph, node size represent the proportion of nodes in the graph while edge width and color represent the proportion of edges between nodes of the respective types. In the figure we omitted node types accounting for less than 1% of the graph nodes.

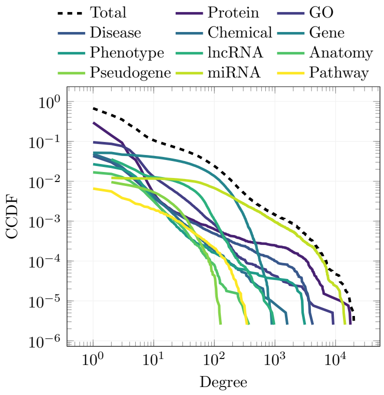

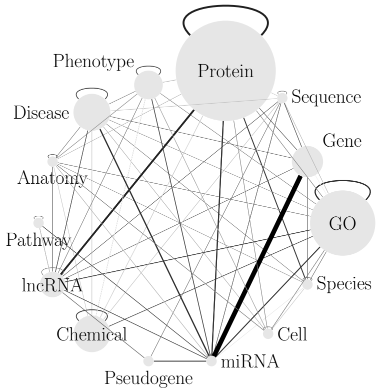

Table S6: Node distribution respect to the node type for the miRNAdisease view*.

| **Node** | **Count** | **Proportion** | **Mean degree** |
|---|---|---|---|
| Protein | 217561 | 0.5524 | 4.0940 |
| GO | 42537 | 0.1080 | 9.6098 |
| Disease | 24073 | 0.0611 | 9.9476 |
| Chemical | 23922 | 0.0607 | 5.3946 |
| Gene | 20376 | 0.0517 | 74.0584 |
| Phenotype | 18713 | 0.0475 | 6.8853 |
| lncRNA | 16641 | 0.0422 | 19.0958 |
| Anatomy | 6597 | 0.0167 | 9.4050 |
| Pseudogene | 5156 | 0.0131 | 13.1389 |
| miRNA | 4800 | 0.0122 | 399.3890 |
| Pathway | 3850 | 0.0098 | 11.4460 |
| Sequence | 2386 | 0.0061 | 24.4778 |
| Species | 2352 | 0.0060 | 42.3899 |
| Cell | 1616 | 0.0041 | 6.5241 |
| Glycan | 797 | 0.0020 | 11.7465 |
| Environment | 446 | 0.0011 | 4.6502 |
| Environmental exposure | 419 | 0.0011 | 3.7542 |

* We report only node types accounting for more than 0.1% of the graph nodes.

Table S7: Edge distribution respect to the type of the edge node extremes for the GOdisease view. We include the most frequent predicates for the extreme node types*.

| Source | Destination | Count | Proportion | Predicates |
|---|---|---|---|---|
| miRNA | Gene | 1439378 | 0.4411 | Regulates activity of (0.3451), Directly negatively regulates activity of (0.2432), Directly positively regulates activity of (0.2427), Indirectly negatively regulates activity of (0.1047), Indirectly positively regulates activity of (0.0643), |
| Protein | Protein | 250975 | 0.0769 | SubClass of (0.9958) |
| Protein | lncRNA | 198287 | 0.0608 | Interacts with |
| miRNA | Disease | 113450 | 0.0348 | Causes or contributes to condition |
| GO | GO | 108502 | 0.0333 | SubClass of (0.8038), Part of (0.0683), Regulates (0.0310), Positively regulates (0.0271), Negatively regulates (0.0269) |
| miRNA | Protein | 81095 | 0.0249 | Interacts with (0.9351), Involved in regulation of (0.0228), Is downstream of sequence of (0.0216), Is upstream of sequence of (0.0201) |
| Protein | miRNA | 79244 | 0.0243 | Interacts with (0.9569), Is upstream of sequence of (0.0221), Is downstream of sequence of (0.0206) |
| Protein | Species | 65636 | 0.0201 | Only in taxon |
| miRNA | Pseudogene | 60011 | 0.0184 | Regulates activity of |
| miRNA | GO | 58472 | 0.0179 | Participates in (0.7063), Has function (0.0874), Interacts with (0.0810), Involved in (0.0498), Located in (0.0428) |
| GO | miRNA | 54571 | 0.0167 | Has participant (0.7568), Function of (0.0937), Interacts with (0.0868), Location of (0.0459), Enabled by (0.0099) |
| Chemical | Chemical | 49896 | 0.0153 | SubClass of (0.7638), Has role (0.2101) |
| Disease | Disease | 43545 | 0.0133 | SubClass of (0.9435), Has characteristic (0.0371), Predisposes towards (0.0086) |
| miRNA | Phenotype | 39220 | 0.0120 | Causes or contributes to condition |
| Phenotype | Phenotype | 37751 | 0.0116 | SubClass of (0.6605), Has modifier (0.1679), Has quality (0.1567), Has part (0.0106) |
| lncRNA | GO | 36476 | 0.0112 | Obsolete contained in (0.7315), Ubiquitously expressed in (0.1131), Located in (0.0639), Under-expressed in (0.0583), Over-expressed in (0.0249) |

* We report only pair of node types accounting for more than 0.1% of the graph edges; and for each pair of node types, the predicates accounting for more than 1% of the edge set.

#### S2.4 piRNAdisease

Figure S6: Left: Complementary Cumulative Distribution Function (CCDF) of the node degrees by node type for the miRNAdisease view. Right: Graph induced by the view types. In the graph, node size represent the proportion of nodes in the graph while edge width and color represent the proportion of edges between nodes of the respective types. In the figure we omitted node types accounting for less than 1% of the graph nodes.

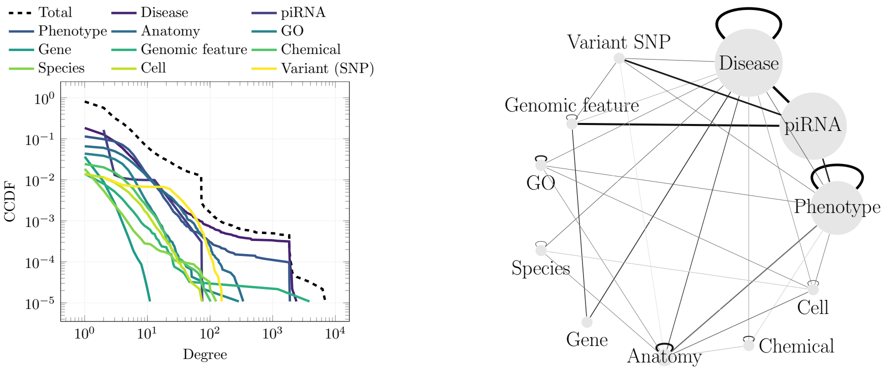

Table S8: Node distribution respect to the node type for the piRNAdisease view*.

| Node | Count | Proportion | Mean degree |
|---|---|---|---|
| Disease | 24203 | 0.2624 | 6.6498 |
| piRNA | 23956 | 0.2597 | 5.3265 |
| Phenotype | 19152 | 0.2076 | 5.5463 |
| Anatomy | 6084 | 0.0660 | 8.6925 |
| GO | 4050 | 0.0439 | 5.9193 |
| Gene | 3823 | 0.0414 | 2.2090 |
| Genomic feature | 2391 | 0.0259 | 14.7633 |
| Chemical | 2239 | 0.0243 | 5.6833 |
| Species | 1784 | 0.0193 | 3.1485 |
| Cell | 1272 | 0.0138 | 5.9057 |
| Variant (SNP) | 1184 | 0.0128 | 23.8657 |
| Environment | 453 | 0.0049 | 4.8212 |
| Environmental exposure | 419 | 0.0045 | 3.7566 |
| Chromosome | 310 | 0.0034 | 5.1839 |
| Protein | 205 | 0.0022 | 7.3366 |
| Food | 181 | 0.0020 | 2.8619 |
| Neuro behaviour | 153 | 0.0017 | 3.7451 |
| Medical action | 114 | 0.0012 | 3.7895 |

* We report only node types accounting for more than 0.1% of the graph nodes.

Table S9: Edge distribution respect to the type of the edge node extremes for the piRNAdisease view. We include the most frequent predicates for the extreme node types*.

| Source | Destination | Count | Proportion | Predicates |
|---|---|---|---|---|
| Disease | Disease | 44117 | 0.1521 | Subclassof (0.9444), Has characteristic (0.0367), Predisposes towards (0.0085) |
| Phenotype | Phenotype | 38489 | 0.1327 | Subclassof (0.6619), Has modifier (0.1672), Has quality (0.1563), Has part (0.0104) |
| piRNA | Disease | 35972 | 0.1240 | Causally influenced by (0.6599), Causes or contributes to condition (0.3401), |
| piRNA | Genomic feature | 23956 | 0.0826 | Subclassof |
| Disease | piRNA | 23738 | 0.0819 | Causally influences |
| Anatomy | Anatomy | 20840 | 0.0719 | Subclassof (0.5189), Part of (0.2486), Develops from (0.0456), Connects (0.0167), Contributes to morphology of (0.0146), Has part (0.0139), Composed primarily of (0.0137) |
| piRNA | Variant (SNP) | 12838 | 0.0443 | Causally influenced by |
| GO | GO | 10978 | 0.0379 | Subclassof (0.7719), Part of (0.0580), Regulates (0.0575), Positively regulates (0.0505), Negatively regulates (0.0504) |
| piRNA | Phenotype | 9130 | 0.0315 | Causally influenced by |
| Phenotype | piRNA | 9130 | 0.0315 | Causally influences |
| Phenotype | Anatomy | 6479 | 0.0223 | Characteristic of (0.5348), Part of (0.2423), Characteristic of part of (0.1375), Occurs in (0.0585), Towards (0.0218) |
| Chemical | Chemical | 5844 | 0.0202 | Subclassof (0.5643), Has role (0.2033), Is conjugate acid of (0.0647), Is conjugate base of (0.0647), Has functional parent (0.0320), Is tautomer of (0.0250), Is enantiomer of (0.0192), Has part (0.0171) |
| Disease | Gene | 4622 | 0.0159 | Has material basis in germline mutation in (0.9929) |
| Gene | Genomic feature | 3823 | 0.0132 | Subclassof |
| Genomic feature | Genomic feature | 3145 | 0.0108 | Subclassof (0.8003), Part of (0.0585), Has quality (0.0525), Has part (0.0483), Derives from (0.0207) |
| Disease | Anatomy | 2996 | 0.0103 | Disease has location (0.8879), Disease has inflammation site (0.0841), Disease arises from alteration in structure (0.0194) |
| Cell | Cell | 2380 | 0.0082 | Subclassof (0.8924), Develops from (0.0983) |
| Species | Species | 1782 | 0.0061 | Subclassof |
| Disease | Phenotype | 1655 | 0.0057 | Disease has feature (0.4961), Has characteristic (0.3994), Disease has major feature (0.0550), Disease arises from feature (0.0387) |
| Disease | Species | 1435 | 0.0049 | In taxon (0.4718), Disease has infectious agent (0.3171), Disease has primary infectious agent (0.0899), Subclassof (0.0669), Transmitted by (0.0523) |
| Variant (SNP) | Genomic feature | 1184 | 0.0041 | Subclassof |
| Environment | Environment | 924 | 0.0032 | Subclassof (0.5260), Part of (0.1082), Composed primarily of (0.0920), Has part (0.0476), Has participant (0.0303), Characteristic of (0.0271), Overlaps (0.0227), Adjacent to (0.0195), Determined by (0.0173), Has output (0.0173), Has input (0.0130), Derives from (0.0119) |
| Variant (SNP) | Disease | 880 | 0.0030 | Causes or contributes to condition |

* We report only pair of node types accounting for more than 0.3% of the graph edges; and for each pair of node types, the predicates accounting for more than 1% of the edge set.

#### S2.5 piRNAdiseaseGO

Figure S7: Left: Complementary Cumulative Distribution Function (CCDF) of the node degrees by node type for the miRNAdisease view. Right: Graph induced by the view types. In the graph, node size represent the proportion of nodes in the graph while edge width and color represent the proportion of edges between nodes of the respective types. In the figure we omitted node types accounting for less than 0.3% of the graph nodes.

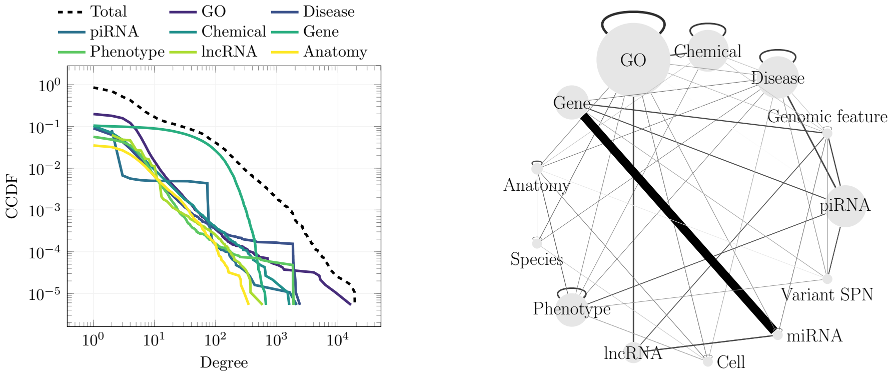

Table S10: Node distribution respect to the node type for the piRNAdiseaseGO view*.

| Node | Count | Proportion | Mean degree |
|---|---|---|---|
| GO | 42385 | 0.2265 | 7.3411 |
| Disease | 24203 | 0.1293 | 6.6499 |
| piRNA | 24048 | 0.1285 | 6.0304 |
| Chemical | 23964 | 0.1281 | 5.4079 |
| Gene | 19591 | 0.1047 | 70.3670 |
| Phenotype | 19168 | 0.1024 | 5.5539 |
| lncRNA | 12105 | 0.0647 | 7.4527 |
| Anatomy | 6561 | 0.0351 | 8.8113 |
| miRNA | 4565 | 0.0244 | 300.6697 |
| Genomic feature | 2391 | 0.0128 | 29.9950 |
| Species | 2370 | 0.0127 | 6.0025 |
| Cell | 1617 | 0.0086 | 6.4917 |
| Variant (SNP) | 1186 | 0.0063 | 23.9553 |
| Protein | 563 | 0.0030 | 5.5240 |
| Environment | 453 | 0.0024 | 4.8234 |
| Environmental exposure | 419 | 0.0022 | 3.7637 |
| Chromosome | 310 | 0.0017 | 5.1839 |
| Plant | 267 | 0.0014 | 4.5543 |
| Food | 181 | 0.0010 | 2.8674 |

* We report only node types accounting for more than 0.1% of the graph nodes.

Table S11: Edge distribution respect to the type of the edge node extremes for the piRNAdiseaseGO view. We include the most frequent predicates for the extreme node types*.

| **Source** | **Destination** | **Count** | **Proportion** | **Predicates** |
|---|---|---|---|---|
| miRNA | Gene | 1333269 | 0.6771 | Regulates activity of |
| GO | GO | 107572 | 0.0546 | Subclassof (0.8036), Part of (0.0681), Regulates (0.0310), Positively regulates (0.0272), Negatively regulates (0.0270) |
| Chemical | Chemical | 50158 | 0.0255 | Subclassof (0.7605), Has role (0.2094), Is conjugate acid of (0.0075), Is conjugate base of (0.0075) |
| Disease | Disease | 44117 | 0.0224 | Subclassof (0.9444), Has characteristic (0.0367), Predisposes towards (0.0085) |
| Phenotype | Phenotype | 38497 | 0.0196 | Subclassof (0.6619), Has modifier (0.1672), Has quality (0.1563), Has part (0.0104) |
| piRNA | Disease | 35972 | 0.0183 | Causally influenced by (0.6599), Causes or contributes to condition (0.3401), |
| GO | Chemical | 28008 | 0.0142 | Has participant (0.7509), Has primary input (0.1467), Has primary output (0.0496), Has primary input or output (0.0445) |
| lncRNA | GO | 27785 | 0.0141 | Located in (0.9962), Involved in (0.0038), |
| GO | lncRNA | 27680 | 0.0141 | Location of |
| piRNA | Genomic feature | 24048 | 0.0122 | Subclassof |
| Disease | piRNA | 23738 | 0.0121 | Causally influences |
| Gene | Genomic feature | 23364 | 0.0119 | Subclassof |
| miRNA | lncRNA | 22644 | 0.0115 | Interacts with |
| lncRNA | miRNA | 22644 | 0.0115 | Interacts with |
| Anatomy | Anatomy | 22502 | 0.0114 | Subclassof (0.5243), Part of (0.2499), Develops from (0.0444), Connects (0.0158), Contributes to morphology of (0.0154), Has part (0.0134), Composed primarily of (0.0129) |
| piRNA | Gene | 17152 | 0.0087 | Regulates activity of |
| piRNA | Variant (SNP) | 12838 | 0.0065 | Causally influenced by |
| Variant (SNP) | piRNA | 12838 | 0.0065 | Causally influences |
| lncRNA | Genomic feature | 12105 | 0.0061 | Subclassof |
| piRNA | Phenotype | 9214 | 0.0047 | Causally influenced by |
| Phenotype | piRNA | 9214 | 0.0047 | Causally influences |
| Phenotype | Anatomy | 6479 | 0.0033 | Characteristic of (0.5305), Part of (0.2423), Characteristic of part of (0.1375), Occurs in (0.0585), Towards (0.0261) |
| GO | Species | 6259 | 0.0032 | In taxon (0.6148), Subclassof (0.3189), Only in taxon (0.0633) |
| miRNA | miRNA | 5750 | 0.0029 | Develops from (0.5000), Develops into (0.5000), |
| Disease | Gene | 4622 | 0.0023 | Has material basis in germline mutation in (0.9924), Disease has basis in dysfunction of (0.0035) |
| miRNA | Genomic feature | 4565 | 0.0023 | Subclassof |
| Species | Species | 3417 | 0.0017 | Subclassof (0.7038), In taxon (0.2962), |
| Genomic feature | Genomic feature | 3145 | 0.0016 | Subclassof (0.8003), Part of (0.0588), Has quality (0.0525), Has part (0.0483), Derives from (0.0207) |
| Cell | Cell | 3005 | 0.0015 | Subclassof (0.8769), Develops from (0.1105) |
| Disease | Anatomy | 2996 | 0.0015 | Disease has location (0.8868), Disease has inflammation site (0.0851), Disease arises from alteration in structure (0.0194) |

* We report only pair of node types accounting for more than 0.1% of the graph edges; and for each pair of node types, the predicates accounting for more than 1% of the edge set.

#### S2.6 chemicaDisease

Figure S8: Left: Complementary Cumulative Distribution Function (CCDF) of the node degrees by node type for the chemicalDisease view. Right: Graph induced by the view types. In the graph, node size represent the proportion of nodes in the graph while edge width and color represent the proportion of edges between nodes of the respective types. In the figure we omitted node types accounting for less than 0.4% of the graph nodes.

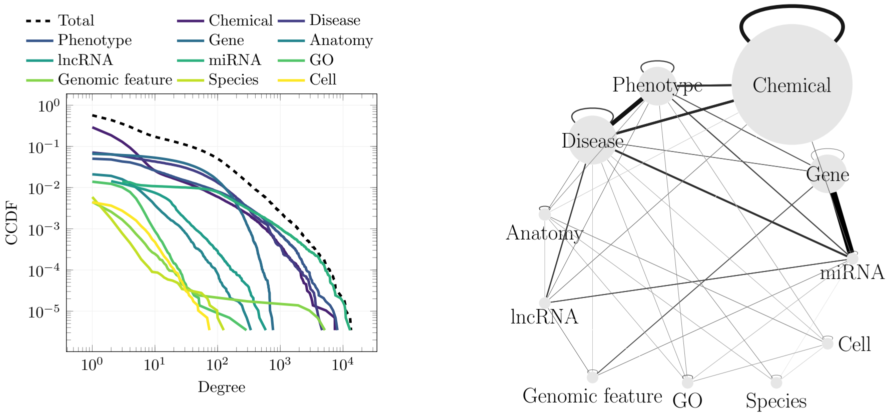

Table S12: Node distribution respect to the node type for the chemicalDisease view*.

| Node | Count | Proportion | Mean degree |
|---|---|---|---|
| Chemical | 199178 | 0.6893 | 6.2203 |
| Disease | 24203 | 0.0838 | 61.0638 |
| Phenotype | 19152 | 0.0663 | 65.7421 |
| Gene | 19075 | 0.0660 | 67.3030 |
| Anatomy | 6084 | 0.0211 | 8.6995 |
| lncRNA | 5123 | 0.0177 | 15.5257 |
| miRNA | 4565 | 0.0158 | 306.4458 |
| GO | 4050 | 0.0140 | 5.9437 |
| Genomic feature | 2391 | 0.0083 | 16.2685 |
| Species | 1784 | 0.0062 | 3.1508 |
| Cell | 1272 | 0.0044 | 5.9033 |
| Environment | 453 | 0.0016 | 4.8256 |
| Environmental exposure | 419 | 0.0014 | 3.7589 |
| Chromosome | 310 | 0.0011 | 5.1839 |

* We report only node types accounting for more than 0.1% of the graph nodes.

Table S13: Edge distribution respect to the type of the edge node extremes for the chemicalDisease view. We include the most frequent predicates for the extreme node types*.

| Source | Destination | Count | Proportion | Predicates |
|---|---|---|---|---|
| miRNA | Gene | 1213775 | 0.3497 | Regulates activity of (0.3696), Directly negatively regulates activity of (0.2426), Directly positively regulates activity of (0.2421) , Indirectly negatively regulates activity of (0.0898), Indirectly positively regulates activity of (0.0558), |
| Disease | Phenotype | 456307 | 0.1315 | Has phenotype (0.9964), Disease has feature (0.0018) |
| Phenotype | Disease | 454652 | 0.1310 | Phenotype of |
| Chemical | Chemical | 368859 | 0.1063 | Subclassof (0.7501), Has role (0.1185), Has functional parent (0.0533), Is conjugate acid of (0.0231), Is conjugate base of (0.0231), Has part (0.0109) |
| Disease | Chemical | 151068 | 0.0435 | Is treated by substance (0.9997) |
| Chemical | Disease | 151019 | 0.0435 | Is substance that treats |
| miRNA | Disease | 105211 | 0.0303 | Causes or contributes to condition |
| Phenotype | Chemical | 99295 | 0.0286 | Is treated by substance (0.9931), Characteristic of (0.0066) |
| Chemical | Phenotype | 98613 | 0.0284 | Is substance that treats Subclassof (0.0000), |
| lncRNA | Disease | 46603 | 0.0134 | Causes or contributes to condition |
| Disease | Disease | 44117 | 0.0127 | Subclassof (0.9444) |
| Phenotype | Phenotype | 38489 | 0.0111 | Subclassof (0.6619), Has modifier (0.1672), Has quality (0.1563), Has part (0.0104) |
| miRNA | Phenotype | 36752 | 0.0106 | Causes or contributes to condition |
| Gene | Phenotype | 24519 | 0.0071 | Causes or contributes to condition |
| Gene | Genomic feature | 22872 | 0.0066 | Subclassof |
| lncRNA | miRNA | 22644 | 0.0065 | Interacts with |
| miRNA | lncRNA | 22644 | 0.0065 | Interacts with |
| Anatomy | Anatomy | 20840 | 0.0060 | Subclassof (0.5189), Part of (0.2517), Develops from (0.0453), Connects (0.0166), Has part (0.0139), Composed primarily of (0.0138), Contributes to morphology of (0.0122) |
| Gene | Disease | 14422 | 0.0042 | Causes or contributes to condition |
| GO | GO | 10978 | 0.0032 | Subclassof (0.7719), Part of (0.0580), Regulates (0.0575), Positively regulates (0.0505), Negatively regulates (0.0504) |
| Phenotype | Anatomy | 6479 | 0.0019 | Characteristic of (0.5309), Part of (0.2423), Characteristic of part of (0.1375), Occurs in (0.0585), Towards (0.0256) |
| miRNA | miRNA | 5750 | 0.0017 | Develops from (0.5000), Develops into (0.5000), |
| lncRNA | Genomic feature | 5123 | 0.0015 | Subclassof |
| Disease | Gene | 4622 | 0.0013 | Has material basis in germline mutation in (0.9937), Disease has basis in dysfunction of (0.0022) |
| miRNA | Genomic feature | 4565 | 0.0013 | Subclassof |
| Chemical | miRNA | 4119 | 0.0012 | Interacts with |
| miRNA | Chemical | 4119 | 0.0012 | Interacts with |
| lncRNA | Phenotype | 3821 | 0.0011 | Causes or contributes to condition |
| Gene | Gene | 3594 | 0.0010 | Genetically interacts with |

* We report only pair of node types accounting for more than 0.1% of the graph edges; and for each pair of node types, the predicates accounting for more than 1% of the edge set.

#### S2.7 cellAnatomy

Figure S9: Left: Complementary Cumulative Distribution Function (CCDF) of the node degrees by node type for the cellAnatomy view. Right: Graph induced by the view types. In the graph, node size represent the proportion of nodes in the graph while edge width and color represent the proportion of edges between nodes of the respective types. In the figure we omitted node types accounting for less than 0.35% of the graph nodes.

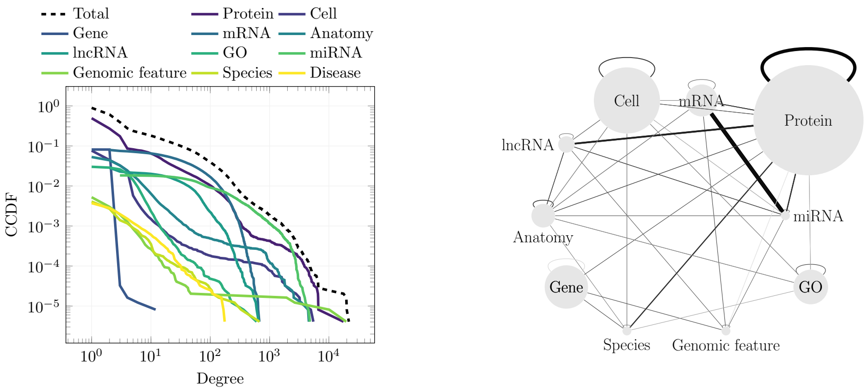

Table S14: Node distribution respect to the node type for the miRNAdisease view*.

| Node | Count | Proportion | Mean degree |
|---|---|---|---|
| Protein | 122682 | 0.4991 | 12.2548 |
| Cell | 40516 | 0.1648 | 4.0871 |
| Gene | 20009 | 0.0814 | 3.0028 |
| mRNA | 19613 | 0.0798 | 63.9203 |
| Anatomy | 14488 | 0.0589 | 12.6432 |
| lncRNA | 10100 | 0.0411 | 28.9183 |
| GO | 7427 | 0.0302 | 6.9841 |
| miRNA | 4565 | 0.0186 | 281.6981 |
| Genomic feature | 2391 | 0.0097 | 25.3371 |
| Species | 1064 | 0.0043 | 114.5385 |
| Disease | 926 | 0.0038 | 7.8197 |
| Chemical | 803 | 0.0033 | 5.6961 |
| Glycan | 797 | 0.0032 | 11.7465 |

* We report only node types accounting for more than 0.1% of the graph nodes.

Table S15: Edge distribution respect to the type of the edge node extremes for the cellAnatomy view. We include the most frequent predicates for the extreme node types*.

| Source | Destination | Count | Proportion | Predicates |
|---|---|---|---|---|
| miRNA | mRNA | 1158937 | 0.3098 | Directly regulates activity of (0.5005), Interacts with (0.4994), Regulates activity of (0.0001), |
| Protein | Protein | 784051 | 0.2096 | Molecularly interacts with (0.7893), Subclassof (0.2093) |
| mRNA | miRNA | 578817 | 0.1547 | Interacts with |
| Protein | lncRNA | 200567 | 0.0536 | Interacts with |
| Protein | Species | 91501 | 0.0245 | Only in taxon |
| miRNA | Protein | 82001 | 0.0219 | Interacts with (0.9361), Involved in regulation of (0.0226), Is downstream of sequence of (0.0214), Is upstream of sequence of (0.0199), |
| Protein | miRNA | 80150 | 0.0214 | Interacts with (0.9578), Is upstream of sequence of (0.0219), Is downstream of sequence of (0.0204), |
| Cell | Cell | 58487 | 0.0156 | Subclassof (0.8577), Derives from (0.1091), Is in cell line repository (0.0279) |
| mRNA | Protein | 45615 | 0.0122 | Interacts with (0.5840), Ribosomally translates to (0.4160), |
| Protein | mRNA | 45615 | 0.0122 | Interacts with (0.5840), Ribosomal translation of (0.4160), |
| Anatomy | Anatomy | 42990 | 0.0115 | Subclassof (0.5582), Part of (0.2571), Develops from (0.0351), Connects (0.0129), Composed primarily of (0.0117) |
| Protein | Protein modification | 32450 | 0.0087 | Has part (0.9999), Subclassof (0.0001), |
| lncRNA | Anatomy | 25679 | 0.0069 | Located in |
| Anatomy | lncRNA | 25679 | 0.0069 | Location of |
| GO | GO | 23719 | 0.0063 | Subclassof (0.7456), Part of (0.0939), Regulates (0.0470), Negatively regulates (0.0407), Positively regulates (0.0404) |
| lncRNA | miRNA | 22644 | 0.0061 | Interacts with |
| miRNA | lncRNA | 22644 | 0.0061 | Interacts with |
| Protein | Gene | 20088 | 0.0054 | Has gene template |
| Gene | Genomic feature | 19976 | 0.0053 | Subclassof |
| Gene | Species | 19975 | 0.0053 | Only in taxon |
| mRNA | Genomic feature | 19613 | 0.0052 | Subclassof |
| Anatomy | Protein | 16214 | 0.0043 | Location of (0.9907), Subclassof (0.0085), Has part (0.0004), Produces (0.0003), Composed primarily of (0.0001), |
| Protein | Anatomy | 16064 | 0.0043 | Located in |
| lncRNA | Genomic feature | 10100 | 0.0027 | Subclassof |
| Cell | Protein | 9862 | 0.0026 | Location of (0.9597), Has plasma membrane part (0.0177), Lacks plasma membrane part (0.0122), Subclassof (0.0044) |
| Protein | Cell | 9473 | 0.0025 | Located in (0.9992), Part of (0.0008), |
| Protein | Glycan | 7769 | 0.0021 | Has part |
| miRNA | miRNA | 7460 | 0.0020 | Develops from (0.3854), Develops into (0.3854), In homology relationship with (0.1147), Genetically interacts with (0.1123) |
| mRNA | mRNA | 7034 | 0.0019 | Interacts with |
| Cell | Species | 6914 | 0.0018 | Part of (0.9954), Only in taxon (0.0022) |
| Cell | Anatomy | 6746 | 0.0018 | Part of (0.9050), Subclassof (0.0434), Derives from (0.0237), Existence starts during (0.0120), Has soma location (0.0099) |
| Cell | Experimental factor | 4731 | 0.0013 | Is disease model for (0.9979), Part of ( |
| miRNA | Genomic feature | 4565 | 0.0012 | Subclassof |
| Cell | Disease | 4318 | 0.0012 | Is disease model for (0.7791), Derives from patient having disease (0.1202), Has disease (0.1003) |

* We report only pair of node types accounting for more than 0.1% of the graph edges; and for each pair of node types, the predicates accounting for more than 1% of the edge set.

### S3 Node Type Prediction

The node-type prediction task aims to predict the type of the $N$ most represented node types. For example, when considering the top 7 node types, the first seven types by frequency (Chemical, Protein, GO, Cell line, Disease, tRF, and mRNA) retain their original labels, while all other types are grouped under a single category.

For this task, embeddings were computed using BFS-like Node2Vec with a skip-gram model (using default GRAPE hyperparameters) and initially tested at the following embedding sizes: 100-dimensional (100-D), 50-D, 10-D, and 2-D. The 2-D embeddings were derived by first generating 100-D~~10-D~~[FT] embeddings and then reducing their dimensionality with t-SNE. These embeddings were subsequently used to train three classification models: decision trees (DTs), random forests (RFs), and radial basis function support vector machines (RBF-SVMs).

The hyperparameters for all models were optimized via 5-fold[FT] stratified grid-search cross-validation on the training set.

Our experiments began with the 2-D embeddings. Supplementary Tables S16, S17, and S18 detail the hyperparameters used for model selection of DTs, RFs, and RBF-SVMs when trained on these t-SNE-reduced embeddings.

Results are shown in supplementary figure S10. RBF-SVMs achieved performance comparable to RFs but required significantly more time to converge. To mitigate this issue, we optimized the RBF-SVM hyperparameters using a subset of the training graph rather than the full dataset. However, the slower convergence and similar performance compared to RFs ultimately led us to exclude RBF-SVMs from further experiments, focusing instead on RFs and DTs.

Next, we optimized DTs (tested hyperparameter values in Supplementary Tables S19) to compare the embedding performance at different embedding sizes (results shown in supplementary figure S11 and supplementary Tables S21 and S22). We determined that 10-D embeddings achieved results comparable to, or slightly better than, the 50-D and 100-D embeddings, which led us to continue our tests with 10-D embeddings.

Supplementary Table S20 presents the hyperparameters used for model selection of RFs when trained on the full 10-D embeddings generated with BFS-like Node2Vec Skip-Gram.

The hyperparameters used to obtain the results reported in the paper for the node-type prediction task are as follows:

- **DT, 2-D embeddings**: Maximum tree depth = 20.
- **DT, 10-D embeddings**: Maximum tree depth = 50, maximum features = 5.
- **DT, 50-D embeddings**: Maximum tree depth = 40, maximum features = 20.
- **DT, 100-D embeddings**: Maximum tree depth = 40, maximum features = 100.

- **RF, 2-D embeddings**: Maximum tree depth = 20, number of trees = 500.
- **RF, 10-D embeddings**: Maximum features = 5, number of trees = 300.
- **SVM, 2-D embeddings**: $C = 315$, $\gamma = 175$.

| Parameter | Values |
|---|---|
| Tree depth | 5, 10, 12, 15, 20, 25, 30, 40, 50 |

Table S16: Hyperparameters and values used for model selection of DTs for node type prediction based on the 2D t-SNE projections of the embeddings.

| Parameter | Values |
|---|---|
| Tree depth | 20 |
| Number of trees | 1, 10, 50, 100, 200, 500 |

Table S17: Hyperparameters and values used for model selection of RFs for node type prediction based on the 2D t-SNE projections of the embeddings.

| Parameter | Values |
|---|---|
| C | 1.0e-02, 1.0e+00, 1.0e+01, 3.2e+02, 1.0e+04, 1.0e+05, 1.0e+07, 3.2e+07, 1.0e+10] |
| $\gamma$ | 1.0e-09, 1.0e-06, 1.0e-03, 1.0e+00, 5.6e+00, 3.2e+01, 1.8e+02, 1.0e+03 |

Table S18: Hyperparameters and values used for model selection of RBF-SVMs for node type prediction based on the 2D t-SNE projections of the embeddings.

### S4 Generic Edge Prediction

The *generic* edge prediction task aims to predict the existence of edges independently of their type. To esnure an unbiased evaluation, we applied a 5-fold stratified cross-validation (70:30, train:test) procedure. Connected Monte-Carlo holdouts were leveraged to select the positive edges of the training set using a spanning tree. The same random seed generator was used to ensure comparability of predictive results. Nodes in the training set were embedded using a BFS-like node2vec approach with Skip-Gram model, employing the default GRAPE hyperparameters and varying embedding sizes (i.e. 10-D, 50-D, and

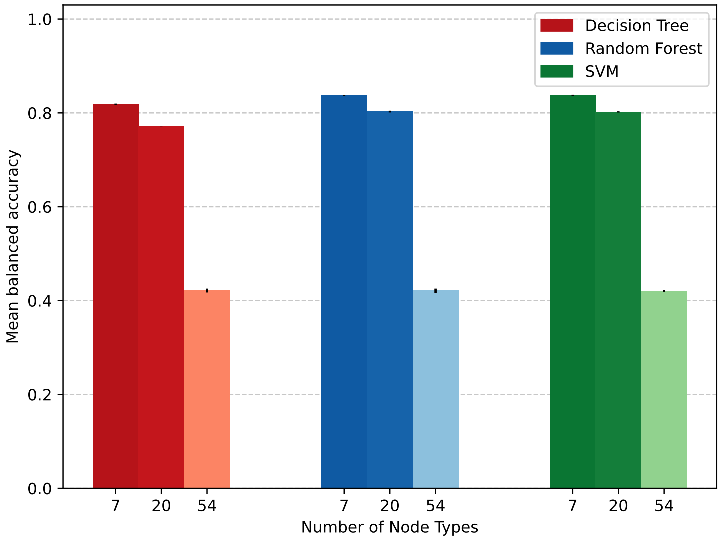

Figure S10: Balanced accuracy results on RNA-KG node type prediction using 2-D embeddings of BFS-like Node2Vec embeddings and DTs (red bars), RFs (blue bars), and RBF-SVM (green bars) as classifier models. Classification results are relative to the 7, 20 and 54 most common node types. The color intensity of the bars reflects their balanced accuracy value. Vertical segments on each point represent the standard deviation across multiple holdouts; note that in some cases the standard deviation is very low ($< 0.1\%$) and the segment is not visible.

| Parameter | Values |
| --- | --- |
| Tree depth | 5, 10, 15, 20, 25, 30, 40, 50, 60, 70, 80 |
| Max features | 1, ‘sqrt’, 5, 7, 9, 10,20,35,40,50,60,80,100 |

Table S19: Hyperparameters and values used for model selection of DTs for node type prediction on 10-D, 50-D, and 100-D embeddings.

| Parameter | Values |
| --- | --- |
| Max features | 3, 5, 7, 10 |
| Number of trees | 50, 100, 150, 200, 250, 300 |

Table S20: Parameters and values used for model selection of RFs for node type prediction based on the full 10-dimensional embeddings.

100-D). The edge embeddings are generated by concatenating the embeddings of the source and destination nodes. Finally, node-degree aware sampling was used to randomly generate negative edges for the training and test sets. Supervised classification was performed using DT and RF models, whose hyperparameters were optimized exploiting a 5-fold stratified grid-search cross-validation on the training set. We considered tree depth, maximum features and number of trees for tuning. Table S23 shows the hyperparameters combinations tested with DT predictors. The number of trees for RF was tuned considering 10, 50, 100, 200 and 500 trees. The other main parameters (tree depth and maximum number of features) were set as the best performing hyperparameters combination found during the DT tuning for each embedding size.

The hyperparameters used to obtain the results shown in the paper for generic edge prediction task are as follows:

- **DT, 10-D embeddings**: 50 maximum tree depth and 10 maximum features.
- **DT, 50-D embeddings**: 200 maximum tree depth and 50 maximum features.
- **DT, 100-D embeddings**: 50 maximum tree depth and 50 maximum features.
- **RF, 10-D embeddings**: 50 maximum tree depth, 10 maximum features, 200 trees.
- **RF, 50-D embeddings**: 200 maximum tree depth, 10 maximum features, 500 trees.
- **RF, 100-D embeddings**: 50 maximum tree depth, 50 maximum features, 500 trees.

| #Classes | Balanced Accuracy | Time (s) |
|---:|---:|---:|
| 7 | 90.29% ± 0.09% | 162.420612 |
| 20 | 88.85% ± 0.12% | 180.897616 |
| 54 | 56.12% ± 1.37% | 331.611162 |
| 68 | 46.82% ± 1.45% | 428.399223 |
| 81 | 46.10% ± 1.97% | 449.408932 |

Table S21: Results obtained with the DT model for node type prediction based on the 50-D embedding with various numbers of node types to predict.

| #Classes | Balanced Accuracy | Time (s) |
|---:|---:|---:|
| 7 | 88.99% ± 0.10% | 857.78 |
| 20 | 87.64% ± 0.09% | 919.63 |
| 54 | 54.60% ± 0.84% | 1185.85 |
| 68 | 44.07% ± 1.41% | 1303.05 |
| 81 | 45.11% ± 2.60% | 1339.91 |

Table S22: Results obtained with the DT model for node type prediction based on the 100-D embedding with various numbers of node types to predict.

| Max features | Tree depth |
|---:|---:|
| 'sqrt' | 10 |
| 'sqrt' | 50 |
| 'sqrt' | 200 |
| 'sqrt' | 500 |
| 25 | 50 |
| 25 | 200 |
| 25 | 500 |
| 50 | 50 |
| 50 | 200 |
| 50 | 500 |

Table S23: Hyperparameters evaluated for the selection of DT models in generic edge type prediction.

### S5 Specific Edge Prediction

The goal of the *specific* edge prediction task is to determine the existence of edges constrained to specific types of source and destination nodes. This task closely aligns with real-world applications, where researchers are often interested in identifying specific novel connections between entities.

To ensure a fair evaluation, we leveraged the unbiased pipeline proposed in Section 3.3 (implementation details in Supplementary Section S6) to con-

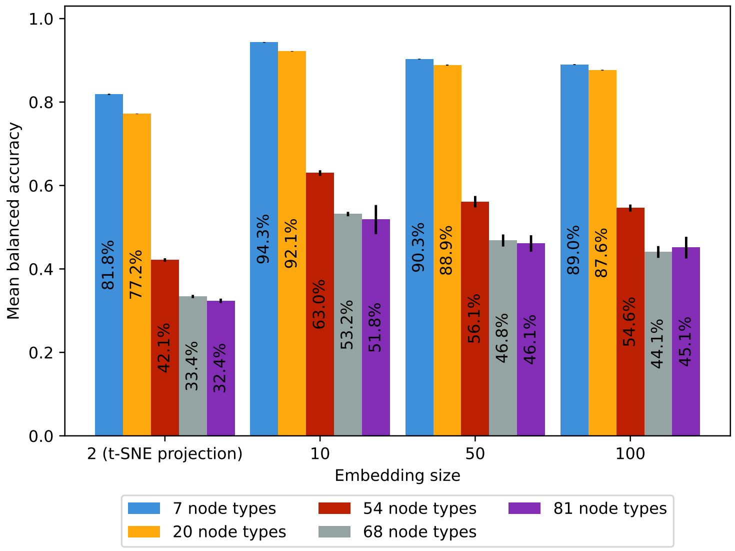

Figure S11: Bar plot comparing the performance of the DT model for node type prediction on the full RNA-KG considering different embedding sizes.

struct positive and negative edges for the training and test sets, considering specific views of RNA-KG (i.e., miRNA-Disease, miRNA-DiseaseSO, GO-Disease, piRNA-Disease, piRNA-DiseaseGO, Chemical-Disease, Cell Anatomy views). Further details about RNA-KG views are available at `https://rna-kg.anacleto.di.unimi.it/views/`.

First-order LINE and BFS-like Node2Vec with Skip-gram model were used to compute the embeddings (size 10-D), which serves as input for supervised classification via DT and RF models. A 5-fold stratified cross-validation (70:30, train:test) was employed to evaluate the generalization performance. The same random seed generator was used to ensure comparability of predictive results.

The hyperparameters used to obtain the results for specific edge prediction are:

- DT: 100 maximum tree depth, 'sqrt' maximum features.
- RF: 100 maximum tree depth and 'sqrt' maximum features and 100 trees.

The following sections report the predictive performance on the RNA-KG views for different types of node pairs.

#### S5.1 MiRNAdisease View

The Balanced Accuracy (BA) results for the specific edge prediction task on the type pairs are reported in table S24 for the LINE embeddings and table S25 for the Node2Vec embeddings.

Table S24: Results of the DT and RF models for specific edge prediction with 10 dimensions embeddings obtained from LINE on miRNAdisease$_{\mathrm{v}}$.

| Model | Type pair | Positive BA | Negative BA | Overall BA |
|---|---|---|---|---|
| DecisionTree | miRNA-Gene | 59.32%±0.14% | 59.34%±0.16% | 59.33%±0.12% |
| DecisionTree | miRNA-Disease | 75.50%±0.36% | 77.04%±0.20% | 76.27%±0.16% |
| DecisionTree | miRNA-Phenotype | 76.16%±0.59% | 77.89%±0.61% | 77.02%±0.39% |
| RandomForest | miRNA-Gene | 65.76%±0.16% | 67.71%±0.12% | 66.73%±0.05% |
| RandomForest | miRNA-Disease | 84.07%±0.20% | 80.80%±0.20% | 82.44%±0.07% |
| RandomForest | miRNA-Phenotype | 87.76%±0.44% | 77.95%±0.23% | 82.85%±0.19% |

Table S25: Results of the DT and RF models for specific edge prediction with 10 dimensions embeddings obtained from Node2Vec Skip-Gram BFS on miRNAdisease$_{\mathrm{v}}$.

| Model | Type pair | Positive BA | Negative BA | Overall BA |
|---|---|---|---|---|
| DecisionTree | miRNA-Gene | 58.60%±0.13% | 56.60%±0.20% | 57.60%±0.12% |
| DecisionTree | miRNA-Disease | 73.89%±0.53% | 76.27%±0.35% | 75.08%±0.24% |
| DecisionTree | miRNA-Phenotype | 75.31%±0.94% | 77.98%±0.75% | 76.65%±0.40% |
| RandomForest | miRNA-Gene | 66.22%±0.17% | 62.18%±0.11% | 64.20%±0.06% |
| RandomForest | miRNA-Disease | 83.58%±0.78% | 80.52%±0.37% | 82.05%±0.21% |
| RandomForest | miRNA-Phenotype | 87.56%±0.31% | 78.14%±0.24% | 82.85%±0.09% |

#### S5.2 MiRNAdiseaseSO View

The Balanced Accuracy (BA) results for the specific edge prediction task on the type pairs are reported in table S26 for LINE and table S27 for Node2Vec.

Table S26: Results of the DT and RF models for specific edge prediction with 10 dimensions embeddings obtained from LINE on miRNAdiseaseSO$_v$.

| Model | Type pair | Positive BA | Negative BA | Overall BA |
|---|---|---|---|---|
| DecisionTree | miRNA-Gene | 59.46%±0.08% | 59.15%±0.14% | 59.30%±0.07% |
| DecisionTree | miRNA-Disease | 75.54%±0.35% | 76.45%±0.25% | 75.99%±0.20% |
| DecisionTree | miRNA-Phenotype | 75.41%±0.44% | 78.86%±0.33% | 77.13%±0.32% |
| RandomForest | miRNA-Gene | 65.98%±0.12% | 67.47%±0.12% | 66.73%±0.05% |
| RandomForest | miRNA-Disease | 84.00%±0.28% | 80.55%±0.26% | 82.27%±0.21% |
| RandomForest | miRNA-Phenotype | 86.41%±0.23% | 79.58%±0.41% | 83.00%±0.24% |

Table S27: Results of the DT and RF models for specific edge prediction with 10 dimensions embeddings obtained from Node2Vec Skip-Gram BFS on miRNAdiseaseSO$_v$.

| Model | Type pair | Positive BA | Negative BA | Overall BA |
|---|---|---|---|---|
| DecisionTree | miRNA-Gene | 58.58%±0.22% | 56.69%±0.29% | 57.63%±0.09% |
| DecisionTree | miRNA-Disease | 74.41%±0.66% | 75.60%±0.28% | 75.00%±0.37% |
| DecisionTree | miRNA-Phenotype | 75.03%±0.91% | 78.64%±0.39% | 76.84%±0.54% |
| RandomForest | miRNA-Gene | 66.01%±0.10% | 62.69%±0.23% | 64.35%±0.09% |
| RandomForest | miRNA-Disease | 83.91%±0.39% | 80.04%±0.22% | 81.98%±0.17% |
| RandomForest | miRNA-Phenotype | 86.48%±0.42% | 79.46%±0.43% | 82.97%±0.31% |

#### S5.3 GOdisease View

The Balanced Accuracy (BA) results for the specific edge prediction task on the type pairs are reported in table S28 for LINE and table S29 for Node2Vec.

#### S5.4 PiRNAdisease View

The Balanced Accuracy (BA) results for the specific edge prediction task on the type pairs are reported in table S30 for LINE and table S31 for Node2Vec.

#### S5.5 PiRNAdiseaseGO View

The Balanced Accuracy (BA) results for the specific edge prediction task on the type pairs are reported in table S32 for LINE and table S33 for Node2Vec.

Table S28: Results of the DT and RF models for specific edge prediction with 10 dimensions embeddings obtained from LINE on $\mathrm{GOdisease_v}$.

| Model | Type pair | Positive BA | Negative BA | Overall BA |
|---|---|---|---|---|
| DecisionTree | miRNA-Phenotype | 75.49%±0.41% | 79.59%±0.32% | 77.54%±0.16% |
| DecisionTree | lncRNA-Phenotype | 58.21%±0.78% | 59.72%±1.71% | 58.97%±0.79% |
| DecisionTree | miRNA-Gene | 67.23%±0.17% | 67.56%±0.14% | 67.40%±0.12% |
| DecisionTree | miRNA-GO | 71.07%±0.43% | 72.21%±0.15% | 71.64%±0.20% |
| DecisionTree | lncRNA-GO | 72.80%±0.98% | 81.54%±0.24% | 77.17%±0.53% |
| DecisionTree | lncRNA-Disease | 65.00%±0.24% | 66.12%±0.67% | 65.56%±0.40% |
| DecisionTree | Protein-GO | 46.37%±4.40% | 61.28%±4.17% | 53.83%±2.09% |
| RandomForest | miRNA-Phenotype | 87.24%±0.30% | 79.37%±0.47% | 83.30%±0.14% |
| RandomForest | lncRNA-Phenotype | 64.11%±0.84% | 65.38%±1.73% | 64.75%±0.61% |
| RandomForest | miRNA-Gene | 72.31%±0.13% | 75.82%±0.10% | 74.07%±0.05% |
| RandomForest | miRNA-GO | 75.32%±0.34% | 81.24%±0.22% | 78.28%±0.25% |
| RandomForest | lncRNA-GO | 79.81%±1.28% | 84.69%±0.33% | 82.25%±0.49% |
| RandomForest | lncRNA-Disease | 73.17%±0.41% | 72.67%±0.22% | 72.92%±0.14% |
| RandomForest | Protein-GO | 36.30%±5.93% | 76.51%±3.46% | 56.41%±1.27% |

Table S29: Results of the DT and RF models for specific edge prediction with 10 dimensions embeddings obtained from Node2Vec Skip-Gram BFS on $\mathrm{GOdisease_v}$.

| Model | Type pair | Positive BA | Negative BA | Overall BA |
|---|---|---|---|---|
| DecisionTree | miRNA-Phenotype | 74.36%±0.46% | 78.92%±0.37% | 76.64%±0.23% |
| DecisionTree | lncRNA-Phenotype | 58.76%±1.68% | 58.51%±1.03% | 58.63%±0.90% |
| DecisionTree | miRNA-Gene | 64.24%±0.80% | 62.53%±1.36% | 63.38%±1.08% |
| DecisionTree | miRNA-GO | 69.45%±0.22% | 68.51%±0.90% | 68.98%±0.51% |
| DecisionTree | lncRNA-GO | 56.83%±2.27% | 88.70%±0.17% | 72.76%±1.10% |
| DecisionTree | lncRNA-Disease | 65.51%±0.74% | 66.32%±0.52% | 65.91%±0.39% |
| DecisionTree | Protein-GO | 45.31%±2.40% | 67.14%±2.83% | 56.22%±1.58% |
| RandomForest | miRNA-Phenotype | 86.60%±0.68% | 79.23%±0.77% | 82.92%±0.25% |
| RandomForest | lncRNA-Phenotype | 64.08%±0.85% | 65.40%±0.87% | 64.74%±0.51% |
| RandomForest | miRNA-Gene | 69.56%±1.09% | 69.00%±2.22% | 69.28%±1.60% |
| RandomForest | miRNA-GO | 73.79%±0.28% | 76.63%±1.95% | 75.21%±0.92% |
| RandomForest | lncRNA-GO | 60.42%±1.90% | 91.25%±0.35% | 75.83%±0.90% |
| RandomForest | lncRNA-Disease | 73.21%±0.74% | 73.76%±0.32% | 73.48%±0.42% |
| RandomForest | Protein-GO | 39.22%±5.70% | 76.54%±4.76% | 57.88%±4.89% |

Table S30: Results of the DT and RF models for specific edge prediction with 10 dimensions embeddings obtained from LINE on piRNAdisease$_v$.

| Model | Type pair | Positive BA | Negative BA | Overall BA |
|---|---|---|---|---|
| DecisionTree | piRNA-Disease | 90.56%±2.74% | 96.13%±0.13% | 93.34%±1.43% |
| DecisionTree | piRNA-Phenotype | N/A[1] | N/A[1] | N/A[1] |
| RandomForest | piRNA-Disease | 89.93%±3.23% | 96.74%±0.13% | 93.34%±1.63% |
| DecisionTree | piRNA-Phenotype | N/A[1] | N/A[1] | N/A[1] |

[1] The data is not available because the model failed to generate a negative training graph with enough edges after the set number of iterations.

Table S31: Results of the DT and RF models for specific edge prediction with 10 dimensions embeddings obtained from Node2Vec Skip-Gram BFS on piRNAdisease$_v$.

| Model | Type pair | Positive BA | Negative BA | Overall BA |
|---|---|---|---|---|
| DecisionTree | piRNA-Disease | 83.05%±3.85% | 99.00%±0.27% | 91.02%±1.79% |
| DecisionTree | piRNA-Phenotype | N/A[1] | N/A[1] | N/A[1] |
| RandomForest | piRNA-Disease | 83.31%±4.11% | 99.78%±0.06% | 91.55%±2.03% |
| RandomForest | piRNA-Phenotype | N/A[1] | N/A[1] | N/A[1] |

[1] The data is not available because the model failed to generate a negative training graph with enough edges after the set number of iterations.

Table S32: Results of the DT and RF models for specific edge prediction with 10 dimensions embeddings obtained from LINE on piRNAdiseaseGO$_v$.

| Model | Type pair | Positive BA | Negative BA | Overall BA |
|---|---|---|---|---|
| DecisionTree | piRNA-Disease | 91.51%±1.86% | 94.90%±0.12% | 93.20%±0.91% |
| DecisionTree | piRNA-Phenotype | N/A[1] | N/A[1] | N/A[1] |
| RandomForest | piRNA-Disease | 91.73%±2.42% | 95.29%±0.34% | 93.51%±1.07% |
| RandomForest | piRNA-Phenotype | N/A[1] | N/A[1] | N/A[1] |

[1] The data is not available because the model failed to generate a negative training graph with enough edges after the set number of iterations.

Table S33: Results of the DT and RF models for specific edge prediction with 10 dimensions embeddings obtained from Node2Vec Skip-Gram BFS on piRNAdiseaseGO$_v$.

| Model | Type pair | Positive BA | Negative BA | Overall BA |
|---|---|---|---|---|
| DecisionTree | piRNA-Disease | 83.17%±3.70% | 98.65%±0.61% | 90.91%±1.54% |
| DecisionTree | piRNA-Phenotype | N/A[1] | N/A[1] | N/A[1] |
| RandomForest | piRNA-Disease | 82.33%±2.69% | 99.62%±0.06% | 90.98%±1.34% |
| RandomForest | piRNA-Phenotype | N/A[1] | N/A[1] | N/A[1] |

[1] The data is not available because the model failed to generate a negative training graph with enough edges after the set number of iterations.

#### S5.6 ChemicalDisease View

The Balanced Accuracy (BA) results for the specific edge prediction task on the type pairs are reported in table S34 for LINE and table S35 for Node2Vec.

Table S34: Results of the DT and RF models for specific edge prediction with 10 dimensions embeddings obtained from LINE on $\text{chemicalDisease}_\text{v}$.

| Model | Type pair | Positive BA | Negative BA | Overall BA |
|---|---|---|---|---|
| DecisionTree | miRNA-Disease | 74.61%±0.36% | 75.66%±0.38% | 75.13%±0.08% |
| DecisionTree | lncRNA-Disease | 72.55%±0.99% | 74.16%±1.13% | 73.36%±1.02% |
| DecisionTree | Gene-Disease | 61.04%±0.50% | 66.65%±0.60% | 63.85%±0.27% |
| DecisionTree | miRNA-Chemical | 58.75%±1.62% | 58.16%±1.23% | 58.45%±0.91% |
| DecisionTree | lncRNA-Chemical | 60.34%±2.65% | 62.29%±2.89% | 61.32%±1.05% |
| DecisionTree | Chemical-Disease | 75.38%±0.23% | 76.40%±0.39% | 75.89%±0.28% |
| RandomForest | miRNA-Disease | 82.87%±0.23% | 79.55%±0.19% | 81.21%±0.03% |
| RandomForest | lncRNA-Disease | 81.36%±1.16% | 79.08%±0.54% | 80.22%±0.63% |
| RandomForest | Gene-Disease | 65.34%±0.99% | 77.02%±0.64% | 71.18%±0.64% |
| RandomForest | miRNA-Chemical | 61.63%±1.46% | 63.19%±0.90% | 62.41%±0.96% |
| RandomForest | lncRNA-Chemical | 71.49%±2.63% | 64.95%±2.99% | 68.22%±1.29% |
| RandomForest | Chemical-Disease | 81.81%±0.09% | 84.64%±0.14% | 83.23%±0.08% |

Table S35: Results of the DT and RF models for specific edge prediction with 10 dimensions embeddings obtained from Node2Vec Skip-Gram BFS on $\text{chemicalDisease}_\text{v}$.

| Model | Type pair | Positive BA | Negative BA | Overall BA |
|---|---|---|---|---|
| DecisionTree | miRNA-Disease | 73.96%±0.45% | 75.54%±0.19% | 74.75%±0.18% |
| DecisionTree | lncRNA-Disease | 68.92%±0.94% | 75.75%±0.47% | 72.33%±0.42% |
| DecisionTree | Gene-Disease | 58.82%±0.34% | 62.47%±0.54% | 60.64%±0.31% |
| DecisionTree | miRNA-Chemical | 58.85%±1.11% | 55.99%±1.50% | 57.42%±0.82% |
| DecisionTree | lncRNA-Chemical | 49.65%±6.37% | 68.64%±3.59% | 59.15%±3.80% |
| DecisionTree | Chemical-Disease | 74.02%±0.22% | 75.57%±0.35% | 74.80%±0.22% |
| RandomForest | miRNA-Disease | 83.31%±0.28% | 80.30%±0.27% | 81.81%±0.08% |
| RandomForest | lncRNA-Disease | 78.68%±0.65% | 81.31%±0.44% | 79.99%±0.26% |
| RandomForest | Gene-Disease | 61.26%±1.11% | 77.66%±0.25% | 69.46%±0.58% |
| RandomForest | miRNA-Chemical | 63.32%±2.32% | 62.27%±0.89% | 62.80%±1.17% |
| RandomForest | lncRNA-Chemical | 51.46%±6.34% | 71.87%±1.87% | 61.66%±2.84% |
| RandomForest | Chemical-Disease | 80.91%±0.22% | 86.49%±0.32% | 83.70%±0.26% |

#### S5.7 CellAnatomy View

The Balanced Accuracy (BA) results for the specific edge prediction task on the type pairs are reported in table S36 for LINE and table S37 for Node2Vec.

Table S36: Results of the DT and RF models for specific edge prediction with 10 dimensions embeddings obtained from LINE on $\text{cellAnatomy}_{\text{v}}$.

| Model | Type pair | Positive BA | Negative BA | Overall BA |
|---|---|---|---|---|
| DecisionTree | Protein-Protein | 77.44%±0.29% | 84.30%±0.28% | 80.87%±0.27% |
| DecisionTree | lncRNA-Anatomy | 79.65%±0.74% | 83.38%±0.69% | 81.51%±0.58% |
| DecisionTree | lncRNA-Cell | 78.78%±2.06% | 85.36%±0.43% | 82.07%±1.15% |
| DecisionTree | Protein-Anatomy | 65.02%±0.75% | 70.47%±0.53% | 67.75%±0.23% |
| DecisionTree | Protein-Cell | 66.93%±1.00% | 70.40%±1.14% | 68.67%±0.56% |
| DecisionTree | lncRNA-Protein | 68.18%±0.22% | 70.29%±0.21% | 69.24%±0.14% |
| DecisionTree | miRNA-Protein | 69.59%±0.26% | 71.46%±0.28% | 70.52%±0.20% |
| RandomForest | Protein-Protein | 79.92%±0.37% | 91.22%±0.08% | 85.57%±0.17% |
| RandomForest | lncRNA-Anatomy | 84.07%±0.49% | 87.87%±0.26% | 85.97%±0.24% |
| RandomForest | lncRNA-Cell | 85.22%±1.13% | 86.50%±0.68% | 85.86%±0.45% |
| RandomForest | Protein-Anatomy | 65.68%±0.27% | 80.13%±0.33% | 72.90%±0.27% |
| RandomForest | Protein-Cell | 69.14%±0.92% | 76.54%±0.73% | 72.84%±0.68% |
| RandomForest | lncRNA-Protein | 77.84%±0.40% | 73.57%±0.37% | 75.71%±0.30% |
| RandomForest | miRNA-Protein | 78.62%±0.36% | 75.30%±0.40% | 76.96%±0.20% |

Table S37: Results of the DT and RF models for specific edge prediction with 10 dimensions embeddings obtained from Node2Vec Skip-Gram BFS on $\text{cellAnatomy}_{\text{v}}$.

| Model | Type pair | Positive BA | Negative BA | Overall BA |
|---|---|---|---|---|
| DecisionTree | Protein-Protein | 71.37%±1.10% | 82.55%±1.27% | 76.96%±1.17% |
| DecisionTree | lncRNA-Anatomy | 78.69%±0.75% | 83.06%±0.73% | 80.88%±0.72% |
| DecisionTree | lncRNA-Cell | 78.13%±1.71% | 86.62%±1.11% | 82.37%±1.17% |
| DecisionTree | Protein-Anatomy | 66.57%±0.26% | 75.41%±1.99% | 70.99%±1.09% |
| DecisionTree | Protein-Cell | 67.34%±2.00% | 72.62%±2.08% | 69.98%±1.95% |
| DecisionTree | lncRNA-Protein | 67.69%±0.32% | 71.68%±0.47% | 69.68%±0.27% |
| DecisionTree | miRNA-Protein | 69.66%±0.31% | 71.59%±0.53% | 70.62%±0.23% |
| RandomForest | Protein-Protein | 76.20%±0.91% | 92.40%±0.59% | 84.30%±0.71% |
| RandomForest | lncRNA-Anatomy | 83.81%±0.23% | 88.05%±1.07% | 85.93%±0.51% |
| RandomForest | lncRNA-Cell | 85.74%±1.27% | 87.59%±0.68% | 86.66%±0.71% |
| RandomForest | Protein-Anatomy | 68.91%±0.75% | 85.83%±1.37% | 77.37%±0.84% |
| RandomForest | Protein-Cell | 71.07%±3.14% | 78.74%±1.68% | 74.91%±2.33% |
| RandomForest | lncRNA-Protein | 76.21%±0.51% | 75.38%±0.60% | 75.80%±0.28% |
| RandomForest | miRNA-Protein | 77.94%±0.40% | 76.15%±0.46% | 77.05%±0.25% |

#### S6 Implementation of the Unbiased Training Procedure

The edge prediction pipeline in GRAPE allows you to pass a positive training set, but it does not allow you to pass a negative training set; instead, the negative training set is generated internally.

According to the code published on the GRAPE GitHub repository, the positive training set used is the one passed as a parameter to the `fit` method. This method handles all the required training operations, so it also takes care of generating the negative training set.

The way the negative training set is generated can significantly impact the model's performance.

According to the repository, the negative training set is generated from the positive training set (`graph` in the code) using the following code:

```
negative_graph = graph.sample_negative_graph(
            number_of_negative_samples =
                number_of_negative_samples,
            only_from_same_component =
                True,
            random_state =
                self._random_state,
            use_scale_free_distribution =
                self._use_scale_free_distribution,
            sample_edge_types =
                len(edge_type_features) > 0,
        )
```

The graph variable contains the positive training graph and `sample_negative_graph` is a method used to generate the negative training set. This method can also accept two additional parameters: `source_node_types_names` and `destination_node_types_names` that are used to generate only some specific edges, but that are not specified in this case, so this method generates a negative training graph that has the same source and destination node types for the edges as the training set.

The main issue with this method is that the graph used to generate the negative training graph is the positive training graph given to the model, which represents only the positive training portion of the holdout set, rather than the full graph. This choice is problematic because some negative edges generated might actually be positive examples that are not included in the training set given to the model but are in the testing set that was left out. This could lead to false negatives that can confuse the predictor being trained, degrading its performance.

#### S6.1 Updating the Training Function

There are two ways to update the training function of a model to change how the training is performed:

- Starting from scratch: this requires reimplementing the model completely, while still ensuring compatibility with GRAPE graphs.
- Updating the classes/objects from GRAPE: this allows for a much faster development time and reduces the risk of errors.

We opted for the second way: updating the classes/objects of the edge prediction models from GRAPE to change the training behaviour.

The GRAPE classes method uses the `_fit` method internally to do the actual training, this method is defined as an abstract method in the abstract class `AbstractClassifierModel`, and it is overridden in the `SklearnLikeEdgePredictionAdapter` class where there is the code to do the actual training.

Figure S12 shows the GRAPE edge prediction models structure for the DT and RF models, along with some possible ways to update it to allow for a custom `_fit` function.

To update the edge prediction classes in GRAPE, we considered three options. The first option (on the right, yellow in figure S12) would be to update the class structure with a custom `SklearnLikeEdgePredictionAdapter` and override the `_fit` function with the custom one at that level to avoid having to create a custom class for each model. This option is difficult to implement because the class is part of a package and updating it does not directly update the subclasses, this would also change the behaviour of the whole edge prediction library branch, which might be unwanted behaviour as well.

The second option (down, blue in figure S12) would be to create a subclass of each edge prediction model we are interested in and override the `fit` function there. This option is easier to implement but requires creating a new class for each model we want to use.

The third option would be to update the instance of the class directly: it is possible to develop a function to update the instance of the edge prediction model, updating the `_fit` function to follow the required behaviour. This option is also easy to implement and does not require creating a subclass for each model, allowing it to be more general.

Both the second and third option were implemented, the outputs produced are the same.

The main changes to apply to the training function are:

1. Add a filter to the negative graph generation function to consider only the specific type pair of interest. This additional step is necessary because the original function used a filtered positive training graph to create the negative graph, while this case uses the original unfiltered graph.
2. Generate the negative graph from the original graph instead of the positive training graph to guarantee no positive edges in the negative training graph.

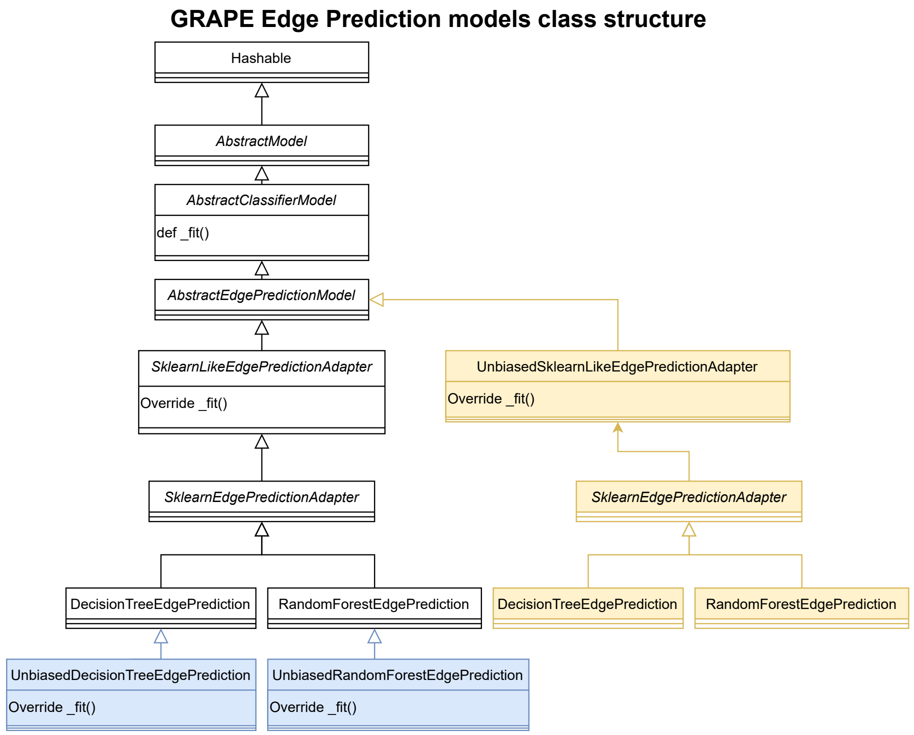

Figure S12: The class structure for the edge prediction models in GRAPE focused on the DT and RF models, with some possible extensions to allow for a custom training function. The classes on the right (yellow) show the option to override the `_fit` function at the general abstract class level, meanwhile the classes at the bottom (blue) show an alternative option, which is to create a custom class that extends each model and overrides the `fit` function as needed.

For the experiments of this paper, the third option was chosen: updating the _fit function. The code is available on GitHub.
